# Bisdemethoxycurcumin mitigates Alzheimer disease pathology through autophagy-mediated reduction of senescence and amyloid beta

**DOI:** 10.1101/2025.05.19.654834

**Authors:** Parul Khajuria, Dilpreet Kour, Kuhu Sharma, Lakhvinder Singh, Razia Banoo, Diksha Manhas, P. Ramajayan, Utpal Nandi, Sandip Bharate, Zabeer Ahmed, Ajay Kumar

## Abstract

AD pathology is accompanied by increased senescence and reduced levels of autophagy in the brain. We investigated whether pharmacologically inducing autophagy could alter the senescent phenotype and help ameliorate AD pathology. We discovered that Bisdemethoxycurcumin (BDMC), a natural compound found in *Curcuma longa*, stimulates autophagy in primary astrocytes. We found that autophagy and senescence exhibit an inverse relationship in aging astrocytes, with increased expression of senescent proteins and downregulation of autophagic proteins. However, treatment of aged astrocytes with BDMC reversed the senescent phenotype by ameliorating the impaired autophagy. Interestingly, the senescent phenotype persisted when autophagy was downregulated by knockdown of *AMPK*. Additionally, BDMC-induced autophagy aided in the removal of amyloid beta that was administered externally to the astrocytes. Further, to validate these results in a mouse model of AD, we confirmed that BDMC can significantly penetrate the blood-brain barrier (BBB) in mice. Therefore, we administered 50 and 100 mg/kg b.w. of BDMC to transgenic 3xTg-AD mice for two months. In their hippocampus, the Control 3xTg-AD animals showed more senescent cells and lower autophagy levels. In contrast, autophagic proteins were significantly upregulated while senescence indicators, such as senescence-associated secretory phenotype (SASP) proteins, were sharply downregulated in the brain of treated animals. Additionally, we discovered that the treated mice’s hippocampus had a significantly lower amyloid beta load. These molecular changes in the brain were ultimately reflected in the improved working memory and neuromuscular coordination behavior of mice treated with BDMC. This study warrants further evaluation of BDMC for the management of AD.

**Graphical Abstract:** 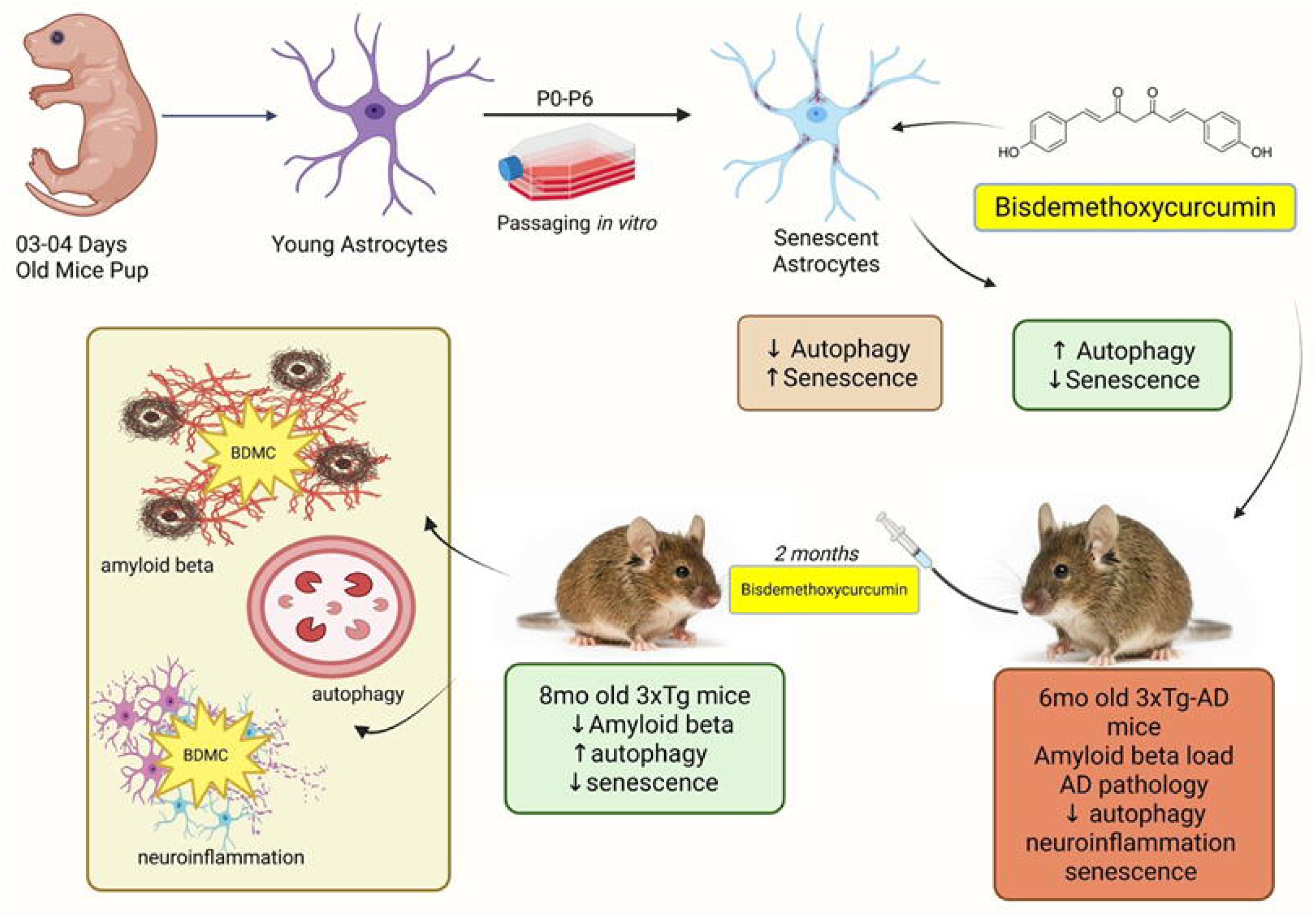
This illustration was created by using biorender.com

## Introduction

Alzheimer’s disease (AD) is the most common form of dementia, characterized by the presence of extracellular neuritic plaques and neurofibrillary tangles (NFTs) and progressive degeneration of the brain tissue^1, 2^. Age-associated functional decline in the physiological processes of the brain is the biggest risk factor for the development of AD ^3^. Physiologically, cells maintain their integrity through a proteolytic mechanism known as autophagy, which helps cells preserve their health by removing protein aggregates and defective organelles like mitochondria, lysosomes, endoplasmic reticulum, etc. However, with age, there is a functional decline in the process of autophagy, leading to the buildup of dysfunctional mitochondria and other organelles, protein aggregates, reactive oxygen species (ROS), and DNA damage ^4^. These cellular defects typically contribute to the onset of senescence in the cells. Thus, impaired or defective autophagy is one of the most important factors that cause cells to enter senescence^5^. Additionally, pharmacological inhibition of autophagy has been strongly linked to the induction of premature senescence ^6^.

Senescence is a natural part of cellular aging, which is accompanied by cell cycle arrest, functional decline and finally death of the cell^7^. During brain ageing, several types of cells, particularly glial cells, show signs of senescence, although several recent studies have implicated senescence in neurons as one of the most important factors responsible for neurodegeneration ^8,9,10^. Senescence in glial cells accelerates the process of neurodegeneration by releasing senescence-associated secretory phenotypes (SASP) such as IL-6 and TNF-α secretion ^10, 11^. AD is a typical example of a neurodegenerative disease where the interplay of autophagy and senescence drives the pathology of the disease. In AD, the decline in autophagy contributes significantly to the accumulation of aggregates of amyloid beta and tau, which act as stimuli for senescence of brain cells ^2, 12^.

In this study, we identified BDMC as a pharmacological inducer of autophagy, which suppresses the onset of senescence to mitigate AD pathology. We aged the primary astrocytes to investigate the interplay of autophagy and senescence in these cells. We proved that a decline in autophagy contributes significantly to the onset of senescence in primary astrocytes. Our data indicated that pharmacological induction of autophagy by BDMC can delay senescence and functional decline of astrocytes. We translated these findings to the in vivo to show that BDMC can cross the BBB. Furthermore, in the transgenic mouse model of AD (3xTg-AD), we showed that it can similarly induce autophagy in the brain and help mitigate AD pathology by delaying senescence and clearing amyloid beta from the hippocampi of mice treated with BDMC.

## Methods and Materials

### Chemicals, Reagents and Antibodies

Chemicals; Dulbecco’s modified eagle medium (DMEM) (Sigma-Aldrich-D1152), Streptomycin (Sigma-Aldrich-S6501), Penicillin (Sigma-Aldrich-P3032), Triton X-100 (Sigma-Aldrich-T8787), Sodium bicarbonate (Sigma-Aldrich-S5761), Phosphate buffer saline (Sigma-Aldrich-D5652), HEPES (Sigma-Aldrich-H3375), Acrylamide (MP biomedical 193982) 4,6-diamidino-2-phenylindole(DAPI) (Sigma-Aldrich-D9542), N,N′-methylene bisacrylamide (Sigma-Aldrich-M7279), Sodium orthovanadate (Sigma-Aldrich-450243), Sodium fluoride (Sigma-Aldrich-215309), Tween 20 (Sigma-Aldrich-P7949), Ammonium persulfate (Sigma-Aldrich-A3678), TEMED (Sigma-Aldrich-T7024),Glycerol (Sigma-Aldrich-G5516), Hanks’ Balanced Salt Solution (HBSS) (Sigma-Aldrich-H6648), Paraformaldehyde (Sigma-Aldrich-P6148), Protease inhibitor cocktail (Sigma-Aldrich-P8340), Trizma-Base (Sigma-Aldrich-T6066), SDS (Sigma-Aldrich-L3771), Fetal Bovine Serum (FBS) (GIBCO-10270106), Acrylamide (MP Biomedical-193982), Glycine (MP Biomedical-194825), Albumin Bovine Fraction V (MP Biomedical-160069), Phenylmethylsulphonyl fluoride (PMSF) (MP Biomedical-195381),Skimmed milk (Himedia-GRM1254), Sodium chloride (Himedia-MB023), Sodium hydroxide(Himedia-MB095),hanks balanced salt solution( Sigma-Aldrich-H6648), Bafilomycin A1 (Sigma-Aldrich-B1793) Rapamycin(Sigma-Aldrich-R8781) Resveratrol(Sigma-Aldrich-R5010) H2DCFDA(Sigma-Aldrich-D6883)Glycerol (Sigma-Aldrich-G5516) MTT(Sigma-Aldrich-M5655) Opti-MEM (gibco-11058-021) beta-amyloid(Anaspec-AS-60479). X-gal (Thermofisher R0404), DMF (Sigma-Aldrich-NSC5356) Potassium Ferrocyanide (Sigma-Aldrich-ATEH99D1EC14) Potassium Ferricyanide (Sigma--Aldrich-702587) AICAR (10010241) (**Antibodies;** AMPK (2532S), pAMPK (2535S), pULK1 (14202S), FIP200 (12436S), MTOR (2972S), pMTOR (5536S),VAMP8 13060, GFAP (80788S), IBA1 (17198S), HRP-linked anti-rabbit IgG (7074S), HRP-linked anti-mouse IgG (7076S), anti-mouse IgG Alexa flour 488 (4408S), anti-mouse IgG Alexa flour 555 (4409S), anti-rabbit IgG Alexa flour 488 (4412S), anti-rabbit IgG Alexa flour 555 (4413S) and siRNA PRKAA2(6620S) were purchased from Cell Signalling Technology. ULK1 (SC-33182), BECN1 (SC-48341), ATG5 (SC-133158), ATG7 (SC-33211), LAMP1 (SC-20011), p53 (sc126), p38 TOMM20 (SC17764) p38 (SC7972) were purchased from Santacruz. Anti-ACTB (A3854), anti-LC3B-II (L7543) and anti-p62/SQSTM1 (P0067) antibodies were purchased from Sigma-Aldrich. Parkin (PA5-13399), p16INK4A (MA5-17054), Cathepsin B (PA5-14255) from Thermofisher. **Kits and other reagents;** PVDF Membrane (Millipore-ISEQ00010), ECL-kit (Millipore-WBKLS0500), Precision plus protein markers (Bio-Rad-161-0375), Bradford reagent (Bio-Rad-500-0006), Mouse IL-1β ELISA kit (Invitrogen-88-7013-88), Mouse IL-6 ELISA kit (Invitrogen-88-7064-88), Mouse TNF-alpha ELISA kit (Invitrogen-88-7324-88) Beta gal senescence green detection kit (2486596) MitoTracker red (M7512), Lysotracker Red (L7528) from Invitrogen.

### Primary astrocytes culture

Culturing of primary astrocytes from mouse pups was performed with approval from the Institutional Animal Ethics Committee (IAEC) of CSIR-IIIM (IAEC approval number: 287/80/2/2022, Study numbers: 0013/2022, 106/2022, 151/2022). Astrocytes were isolated from the brains of 3-4-day-old C57BL/6J mice pups and were grown in 10% FBS-supplemented DMEM media. Briefly, pups were sacrificed in a CO2 chamber, and the brain was dissected out. The cortical region separated using a fine blade was collected in incomplete media and cut into small pieces. After that, these cortices were incubated at 37°C in 0.25 % trypsin for 30 minutes, with shaking after every 10 minutes. Trypsin activity was neutralized with complete media, and cells were collected using a 70-micron cell strainer followed by centrifugation at 300 rpm for 5 minutes. Pellet was dissolved in fresh media, and cells were grown in cell culture flasks and were maintained at 37°C in a CO_2_ incubator. Microglia and oligodendrocytes were separated from the underlying astrocytes layer after 6-7 days by shaking the flask at 240 rpm for 6 hours.

The sub-cultured astrocytes were assessed for enrichment of culture through glial fibrillary acidic protein (GFAP) marker, and purity was found to be more than 95% based on the GFAP marker and markers for microglial cells (IBA1), and neuronal cells (NEUN) (Supplementary Figure S1A). The first sub-cultured astrocytes were used as passage 1 (P1) and were incubated until confluent growth. After that, P1 astrocytes were sub-cultured into 2.5×10^5^ cells/ml to make the second passage (P2). These processes were repeated until passage 6 (P6) to make the late passage astrocytes for carrying out the experiments.

### Extraction and isolation of Bisdemethoxycurcumin (BDMC)

One kg of dried and powdered rhizomes of *Curcuma longa* was extracted using a Soxhlet apparatus with 3 L of methanol. The methanol was evaporated under reduced pressure using a rotary evaporator, yielding an orange residue. This residue was purified through silica gel column chromatography with a dichloromethane/methanol eluent to isolate Bisdemethoxycurcumin. The product was characterized by NMR and MS analyses. 1H NMR (400 MHz, DMSO-d6) δ 10.05 (s, 1H, enol OH), 7.58 (d, J = 8.6 Hz, 4H), 7.52 (d, J = 12 Hz, 1H), 6.83 (d, J = 8.6 Hz, 4H), 6.70 (d, J = 15.9 Hz, 2H), 6.04 (s, 1H), 4.05 (s, 2H).13C NMR (101 MHz, DMSO-d6) δ 188.39, 165.01, 145.58, 135.53, 131.04, 125.96, 121.15, 106.25. ESI-MS (-ve): m/z 307 [M-H].+ (The NMR spectra scans are provided in Supplementary Information, Figure S1B and S1C).

### Autophagy flux measurement

For measuring autophagy flux, astrocytes at P2 were treated with 2.5 µM BDMC for 24 h, and astrocytes from P4-P6 were treated with 2.5 µM BDMC for 24 h at P4, P5, and P6 intervention following the confluency. Cells were also treated with or without the autophagy inhibitor, bafilomycin A_1_. Bafilomycin A_1_ was added 3 h prior to the termination of the experiment at a 20 nM concentration. Levels of LC3B-II and SQSTM1 proteins were measured in cell lysates using an immunoblotting assay.

### Immunoblotting

Cellular lysates were prepared using radioimmunoprecipitation assay (RIPA) buffer for protein analysis. Briefly, the cell pellet was dissolved in RIPA buffer containing sodium fluoride, sodium orthovanadate, phenyl methyl sulfonyl fluoride (PMSF) and 1% protease inhibitor cocktail for 1 h with vortexing sample every 15 min. The supernatant was collected after high-speed centrifugation of samples for 20 min at 4°C. Total protein was measured using the Bradford assay. Proteins were separated using SDS-PAGE run for 3h at 75 V.The electrophoresed proteins were then transferred from the gel to PVDF membrane for 120 minutes at 100 V. The protein of interest was labelled with a specific primary antibody overnight at 4°C. Further, the blot was incubated with corresponding HRP-conjugated secondary antibody for 1 h at room temperature. The band were visualized in the GenaxyChemiDoc imaging system (Make: Syngene, Maryland, USA; Model: G: BOX, XT-4). Densitometry analysis of all the Immunoblots was performed by using Image J software. Band densities of proteins were normalized with β-actin. The phosphorylated proteins were normalized with their total forms.

### Senescence-associated-β-galactosidase (SA-β-gal) staining

For β-gal staining, aged (P6) and young astrocytes (P2) were seeded in 6-well plates and treated with BDMC (2.5 µM) for 24 h. Cells were then washed with PBS and fixed with 4% paraformaldehyde for 5 minutes and incubated in staining solution (0.2 M sodium phosphate buffer, 100 mM potassium ferricyanide and ferrocyanide, 5 M NaCl, 1 M MgCl_2_ and 50 mg/ml X-gal in DMF) at 37 °C overnight. The following day, the cells were washed with PBS and analyzed under a light microscope.

Intracellular senescence-associated β-galactosidase (SA-β-Gal) activity was assayed using an SA-β-Gal staining kit (CellEvent™ Senescence Green Detection Kit) according to the manufacturer’s instructions, and senescent cells were identified as green-stained cells under a high-throughput microscope. The images were acquired and quantified using the Confocal Quantitative Image Cytometer at 20X (Yokogawa CQ1).

### Enzyme-linked Immunosorbent Assay (ELISA)

For detection of SASPs, TNF-α and IL-6, primary astrocytes, both aged and young, were seeded in 6 well plate and treated with BDMC (2.5 µM). The levels of SASPs were measured in the supernatant and analyzed using ELISA following the manufacturer’s guidelines (Invitrogen).

### ROS detection

To detect ROS production, the 2’,7-dichlorofluorescin diacetate (DCFH-DA) (Sigma, D6883) was used. One day prior to the treatments, 2 × 10^5^ cells were plated per wellin a six-well dish. To measure ROS, DCFH-DA (final concentration 10 μM) was added to the media for the last 30 min. After the incubation media was removed, cells were washed once with PBS and fluorescence intensity was measured using a spectrophotometer at excitation 498 nm and emission 522 nm. Analysis was performed on a Tecan spectrophotometer microplate reader.

### Confocal microscopy

Astrocytes were seeded on coverslips in 6-well plates. After treatment with BDMC (2.5 µM) for 24 h, cells were fixed with 4% paraformaldehyde (PFA) for 15 min and permeabilized with 0.1% Triton-X 100 for 7 min. Blocking was done for 30 min with 2% BSA, followed by overnight incubation with the primary antibody. Alexa flour-conjugated secondary antibody incubation was given to cells for 1 h for labelling. The nucleus was stained with 4,6-diamidino-2-phenylindole (DAPI) (1µg/ml) for 10 min. Following staining, the coverslips were mounted on slides using mounting media (glycerol and PBS at a ratio of 9:1). Cells were washed thrice with PBS after each incubation. The images were acquired in a CQ1 high-throughput imaging system.

### Aβ clearance assay

Aβ_42_-HiLyte flour488 peptide was prepared following the manufacturer’s protocol (Anaspec Inc.). Primary astrocytes seeded on coverslips were treated with BDMC (2.5 µM) or Rapamycin (250 nM). After 12 h, 2 µg/ml fluorochrome-tagged (Hilyte flour488) Aβ_42_ proteins were added to cells for another 12 h. Bafilomycin A_1_ was given 3 h prior to experiment termination. After the end of the assay, cells were washed with PBS and cells were fixed with 4% PFA, permeabilized with Triton-X 100, and the nucleus was stained with DAPI, as mentioned above. Similarly, slides were prepared, and images were taken and analyzed in the CQ1 high-throughput imaging system.

### Transfection of astrocytes with *siAMPK*

AMPK expression was knocked down in astrocytes by using siRNA against AMPK using FuGENE HD for 24 h. Primary astrocytes were seeded in 6-well plates at a density of 0.4 × 10^6. The cell media was replaced by OPTI-MEM media for 30 min prior to transfection. The transfection was done using Lipofectamine 3000 (Invitrogen) in the ratio 1:200. The transfection mixture was prepared and kept in sterile conditions for 20 min before being added to the cells. Following this, cells were assessed for autophagy and senescence through confocal microscopy and western blotting.

### Pharmacokinetic analysis

MaleC57BL/6J mice having a body weight of 25-30 g were used for the plasma and brain pharmacokinetic study of Bisdemethoxycurcumin (BDMC). Animals were maintained in standard laboratory conditions, i.e., room temperature (25 ± 2°C), light/dark cycle (12 h/12 h), and the relative humidity (50 ± 20 %), with free access to standard pellet diet and water *ad libitum*. On the day of the experiment, mice were randomly divided into six groups containing five animals each for sparse sampling. The study was performed following a single dose oral administration of BDMC at 50mg/kg body weight. Formulation was prepared by using the vehicle:5% DMSO + 20% Tween-80 + 45% PEG-400 + q.s. Water (v/v). The dose volume was used at 10 ml/kg. Blood and brain samples were collected at pre-determined time points such as 0, 0.25, 0.5, 1, 2, and 4 h. Blood samples were collected from the retro-orbital plexus into the micro centrifuge tubes containing 5% (w/v) disodium EDTA as an anticoagulant, followed by brain extraction. For plasma sample analysis, each plasma sample was centrifuged at 14000 rpm for 10 min to obtain 50 µL of plasma and stored at -80°C till analysis. On the day of analysis, the plasma sample was thawed and processed by the addition of 200 µl of ethyl acetate containing IS (50 ng/mL) for plasma protein precipitation. After that, the sample was vortexed thoroughly for 2 min, followed by centrifugation at 14000 rpm for 10 min at 4°C. Then, the solvent layer was decanted and dried under vacuum. After this, the sample was reconstituted with 150 µL of acetonitrile and analyzed by LC-MS/MS [SRM transition of BDMC and IS (Diazepam) was 309.1 > 225.1 and 285.1 > 193.0 in Positive mode]. For brain sample analysis, brain homogenate was prepared in PBS (10 mM, pH 7.4) at 350 mg/ml. The brain homogenate (50 µL) sample was then processed and analyzed as mentioned above for plasma samples.

### Animals and Ethical Clearance

Six-month-old 3xTg-AD transgenic mice were used in the study and randomly divided into three groups (n = 7). C57BL/6J mice were used as Control wild type (WT). They were housed under a 12 h light/dark cycle in a temperature (65–75 °F; 18-23 °C) and humidity-controlled (40–60 %) environment, supplied with free access to food and water (ad libitum). Prior to the initiation of the study, all the animals were acclimatized for one week under standard laboratory conditions. The animals were drug naive with no prior procedures performed. Mice were randomized into groups based on their body weights for all behavioral assays. The total study duration was 2 months. All experiment protocols were approved by the Institutional Animal Ethics Committee (IAEC) (IAEC approval no. 321/82/2/2023, study no. 52/82/2/2023) and followed the Committee for Control and Supervision of Experiments on Animals (CCSEA; Ministry of Environment and Forest, Government of India) guidelines for animal care.

### Behavioral analysis

To check the effect of BDMC on the behavior of 3xTg-AD mice, open field test (OFT), rotarod (Make; Ugo Basile, Accelerrota-rod for mice 7650) and radial arm maze (RAM) tests were performed.

### Rota rod test

To check the effect of BDMC on neuromuscular coordination, the rotarod test was performed. After training mice to run on the rotarod for five days, their latency to fall was calculated on the day of analysis.The rotarod was initially set at a speed of 4 rpm at the start of the test and accelerated to 40 rpm over 300 s. The latency to fall from the rotarod was recorded for each mouse, and the mean latency to fall for each group was calculated

### Open field test

OFT was performed to assess exploratory and locomotor activity in a white box (45 × 60 × 25 cm). Distance travelled, speed, time spent in corner and center zones and mobile episodes were recorded through an automated video tracking system using AnyMaze software.

### Radial arm maze

To check the effect of BDMC on the spatial memory of mice, a radial arm maze test was performed on an eight-armed radial arm maze (UGO Basile). Mice were trained for five days to locate baited arms with visual clues, and on the day of analysis, several parameters were checked using AnyMaze software. On the memory retention trial day, the mouse was placed at the end of one arm (entry arm) and allowed to navigate the baited arm. The movement of mice was tracked with the help of automated AnyMaze software connected with a video tracking camera.

### Immunohistochemistry of 3xTg-AD brain

After the treatment completion, the hippocampi were isolated from the mice brains and the 10 μm-thick tissue sections were prepared. For IHC, the slides were deparaffinized in xylene and rehydrated through an ethanol dilution series (100-95-70-50%). Endogenous peroxidase activity was blocked by incubating sections in 3% H_2_O_2_ for 15 min. For immunostaining of Aβ_42_ and GFAP, the sections were pre-treated with microwave in citrate buffer pH 6.0 for 20-25 minutes for epitope retrieval.

Tissues were incubated with primary antibodies for 12 h, followed by HRP-conjugated anti-rabbit, which was used as the secondary antibody. Freshly prepared DAB substrate solution was added to slides for minutes. The slides were counterstained with hematoxylin, dehydrated in ethanol (50-70-95-100%), cleared in xylene and mounted on mounting media (Sigma). Staining was evaluated under the light microscope.

### Western blot analysis of proteins in the mouse hippocampi

The hippocampi isolated from mice brains were immediately washed with saline and stored in liquid nitrogen. For lysate preparation, tissue was homogenized in RIPA buffer substituted with sodium fluoride, sodium orthovanadate, phenyl methyl sulfonyl fluoride and 1% protease inhibitor cocktail. Total protein was measured by Bradford assay. The isolated protein supernatant was diluted with 2X Lammeli buffer and subjected to western blot assay as described above for protein analysis.

### Detection of SASPs (senescence-associated secretory phenotypes) in the mouse cortex

Mouse cortices were isolated and homogenized in RIPA buffer. The levels of different SASPs were analyzed by performing ELISA on the homogenates following the manufacturer’s protocol. (Invitrogen)

### Statistical Analysis

Statistical analyses were performed using GraphPad Prism 8 software. All in-vitro experiments were performed thrice. The response of independent mice from the group was noted for animal experiments, and the mean was calculated. The data presented here represents the mean[±[SD. Statistical significance of the experiment was calculated using one-way ANOVA analysis, and the Bonferroni method was used as a post-hoc test. p value less than 0.05 was taken as significant with values assigned as ****p < 0.0001, ***p < 0.001, **p< 0.01, *p< 0.05 and ns = not significant.

## Results

### The aging of primary astrocytes is accompanied by reduced basal level of autophagy and elevated senescence

Astrocytes play a crucial role in maintaining the homeostasis of the brain. We wanted to know if the aging of astrocytes can alter autophagy, a key homeostatic mechanism. Therefore, we aged the primary mice astrocytes up to passage six (P-6) and compared their levels of autophagic proteins with passage two (P-2). We found that with aging, astrocytes display a significant downregulation of pAMPK (Thr 172), the most important regulator of autophagy (Figure 1A). Further investigation revealed an overall slowdown of autophagy, which was reflected through decreased expression of ATG5, ATG7, BECN1, and LC3B-II and increased expression of SQSTM1 (Figure 1A and Figure 1C). We further asked if aging can also affect the levels of senescence in primary astrocytes. Therefore, we compared the P-2 and P-6 astrocytes for the presence of senescence. We found a significant rise in the number of senescent cells with the increasing number of passages of primary astrocytes (Figure 1B and 1D). This increased senescence with aging was analyzed by using molecular markers like p16INK4A, p38, and p53, which showed a marked increase in the expression of these genes, indicating induction of senescence with aging (Figure 1B&D).

**Figure 1:**
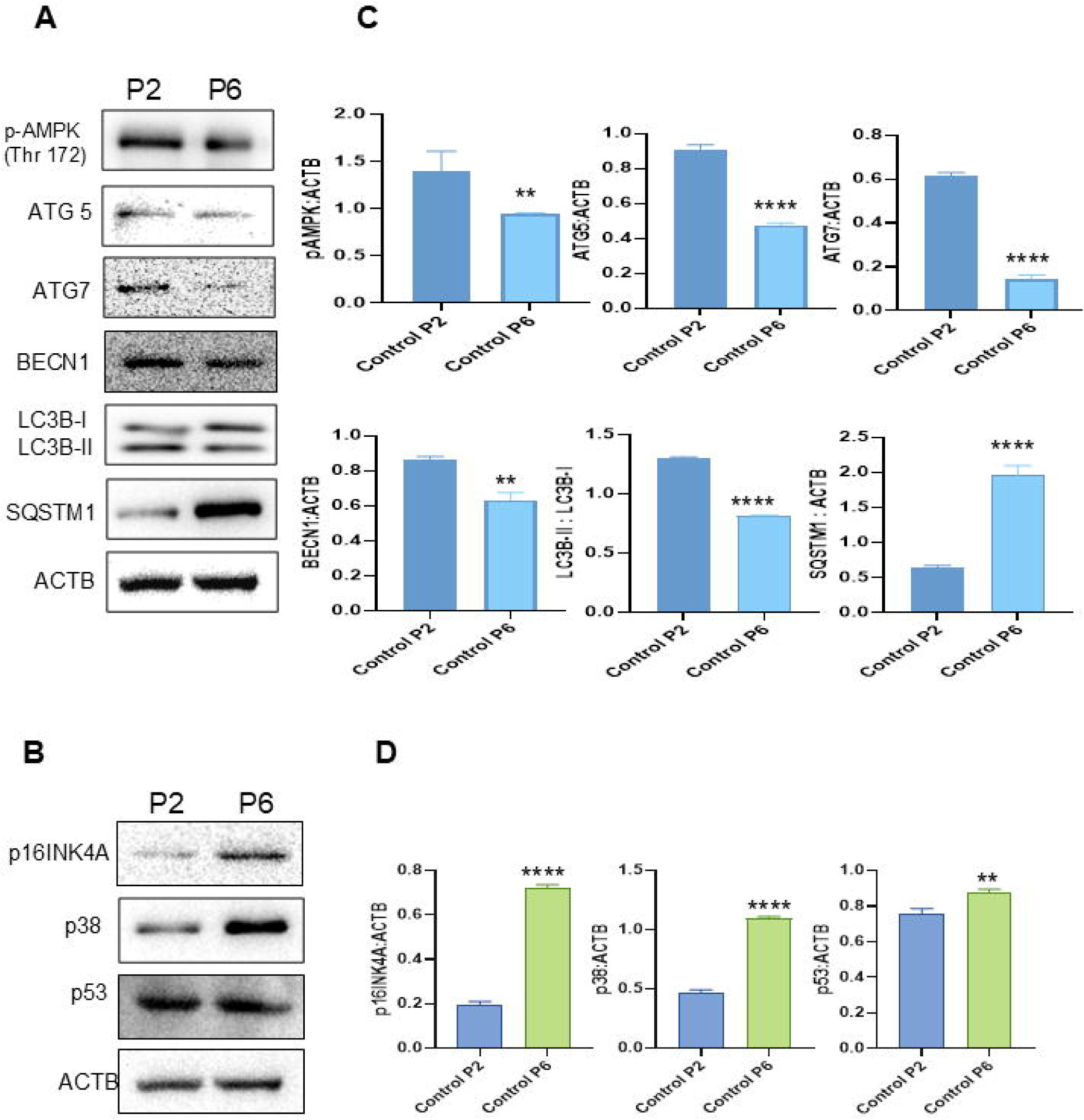
Autophagy is defective and senescence is upregulated in aged primary astrocytes. Primary astrocytes isolated from the brain of 3 to 4 days old C-57 BL6/J mice pups were cultured uptill passage 6 and analyzed.**(A)**Immunoblots depicting down regulation of autophagy pathway proteins.**(B)**Representative western blot images showing upregulated levels of p16, p38and p53, senescence associated pathway proteins in aged astrocytes**. (C)** Quantification of the autophagy protein levels expression normalized with ACTB and measured using the imageJ software. **(D)** Level of senescence associated proteins normalized with ACTB. Statistical significance was analyzed using one way ANOVA and Bonferroni test for multiple comparisons. p values ****p <0.0001,***p < 0.001.

### BDMC upregulates autophagy pathway in primary astrocytes

After establishing that autophagy, a major homeostatic mechanism, is compromised in aged astrocytes, we proceeded to pharmacologically intervene with an effective autophagy inducer, which could alleviate these defects. We found BDMC (Bisdemethoxycurcumin) to be a potent natural compound that could enhance autophagy. BDMC is a naturally occurring curcuminoid isolated from the *Curcuma longa* rhizome^13^. It is a demethoxy derivative of curcumin (Figure 2A). To examine the effect of BDMC on autophagy, we treated primary astrocytes with BDMC for 24 h and assessed the levels of autophagy marker proteins, LC3B-II and SQSTM1 (Figure 2B). Our results showed that BDMC was able to enhance autophagy at a low concentration of 1.25 µM as indicated by elevated levels of LC3B-II, an important marker for autophagy and reduced levels of SQSTM1 (sequestosome1), a ubiquitin-binding adaptor of the autophagy pathway (Figure 2B-C). These results were further validated by the effect of BDMC on levels of various other autophagy-related proteins. We found that BDMC elevated the levels of key autophagy pathway proteins, pAMPK (Thr172), FIP200, pULK1, BECN1, ATG5, ATG7, VAMP8 and CTSB and reduced the mTOR (Ser4228) levels, which act as an inhibitor of autophagy (Figure 2D and Supplementary Figure S2A). Based on the concentration-dependent activity of BDMC and its ability to enhance autophagy, we selected 2.5 µM as a working concentration for further studies (Figure 2B-D and Supplementary Figure S2A).

**Figure 2:**
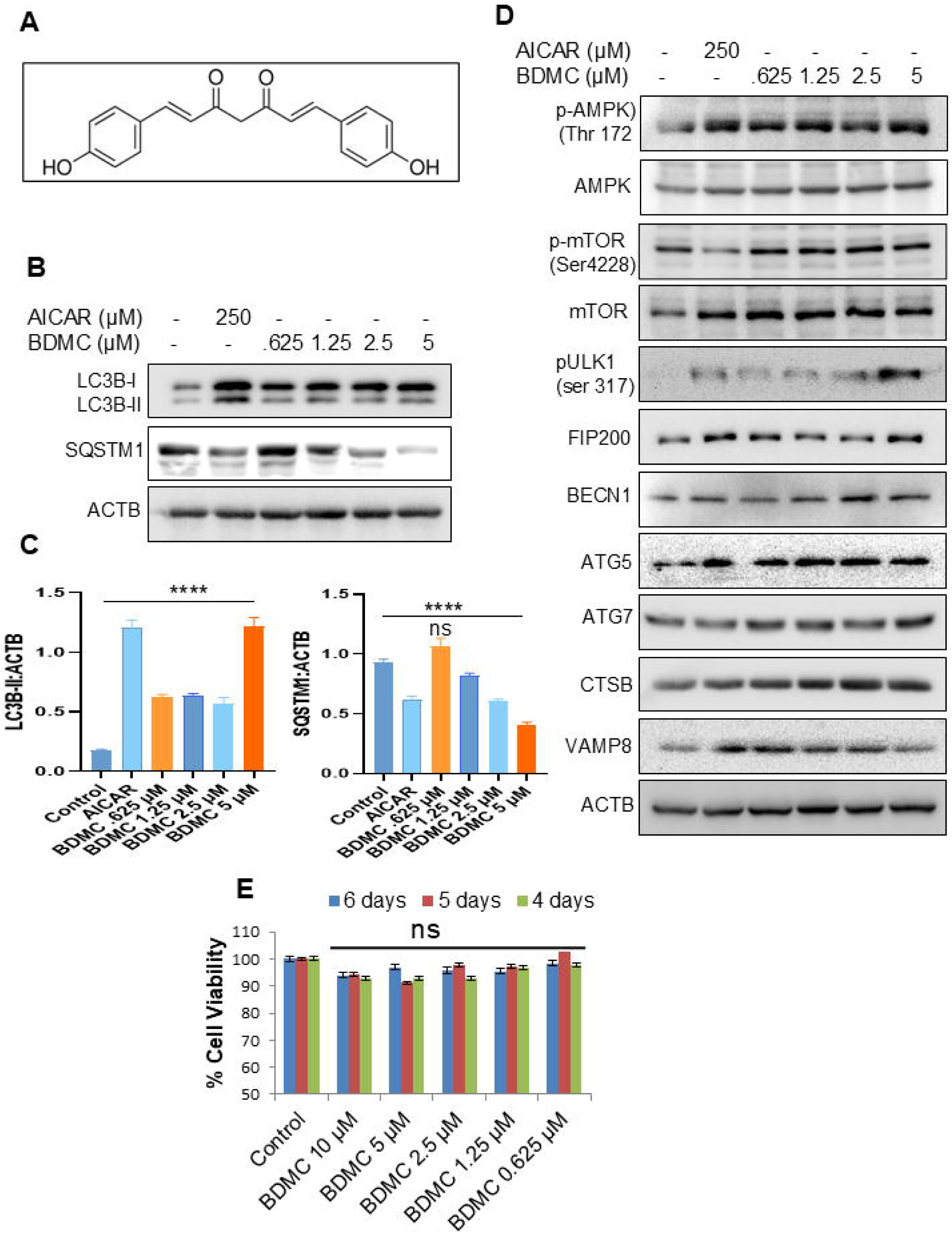
BDMC induces autophagy in primary astrocytes. **(A)** Structure of BDMC. **(B)** Western blot analysis of key autophagy proteins LC3B-II and SQSTM1 after treatment with BDMC (2.5 µM). **(C)** Quantification of western blots shown in fig (B) **(D)** Immunoblots of autophagy pathway proteins in primary astrocytes after treatment with BDMC for 24 h and AICAR (250 µM) as standard autophagy inducer for 3h. **(E)** Safety profiling of BDMC measured through MTT assay following 24h

Autophagy as a homeostasis mechanism is dysregulated with aging and age-related neurodegenerative disorders, particularly AD, which leads to the accumulation of protein aggregates. To evaluate the effect of BDMC on defective autophagy in aged astrocytes, we first needed to check whether BDMC was safe for astrocytes when administered for a prolonged period of time. For this, primary astrocytes were incubated with various concentrations of BDMC for 4, 5 and 6 days, respectively. Through MTT assay, we found that astrocytes remained viable even after prolonged treatments with BDMC (Figure 2E and Supplementary Figure S2B), and BDMC had no adverse effect on cell viability. Taken together, these findings suggest that BDMC is a viable and potent agent capable of inducing autophagy at a low concentration.

### BDMC restores autophagy and prevents senescence in aged astrocytes

Defective autophagy in neurodegenerative diseases like AD (Alzheimer disease) is associated with an increased incidence of senescent phenotype in the brain^14^ . To investigate whether prolonged treatment with BDMC could reduce senescence with age and alleviate dysregulated autophagy, we aged primary astrocytes and at P4, P5 and P6 intervention with 2.5 µM BDMC was given. Given that a decline in autophagy flux is associated with defective autophagy in AD, we found that BDMC could enhance the levels of autophagy flux in aged primary astrocytes (Figure 3A and supplementary Figure S3A). These senescent astrocytes treated with bafilomycin A1 showed a higher blockade of autophagy flux, indicating impairment in lysosome autophagosome fusion. However, with BDMC treatment, levels of LC3II-B increased and SQSTM1 decreased when analyzed through western blotting (Figure 3A and Supplementary Figure S3A).

**Figure 3:**
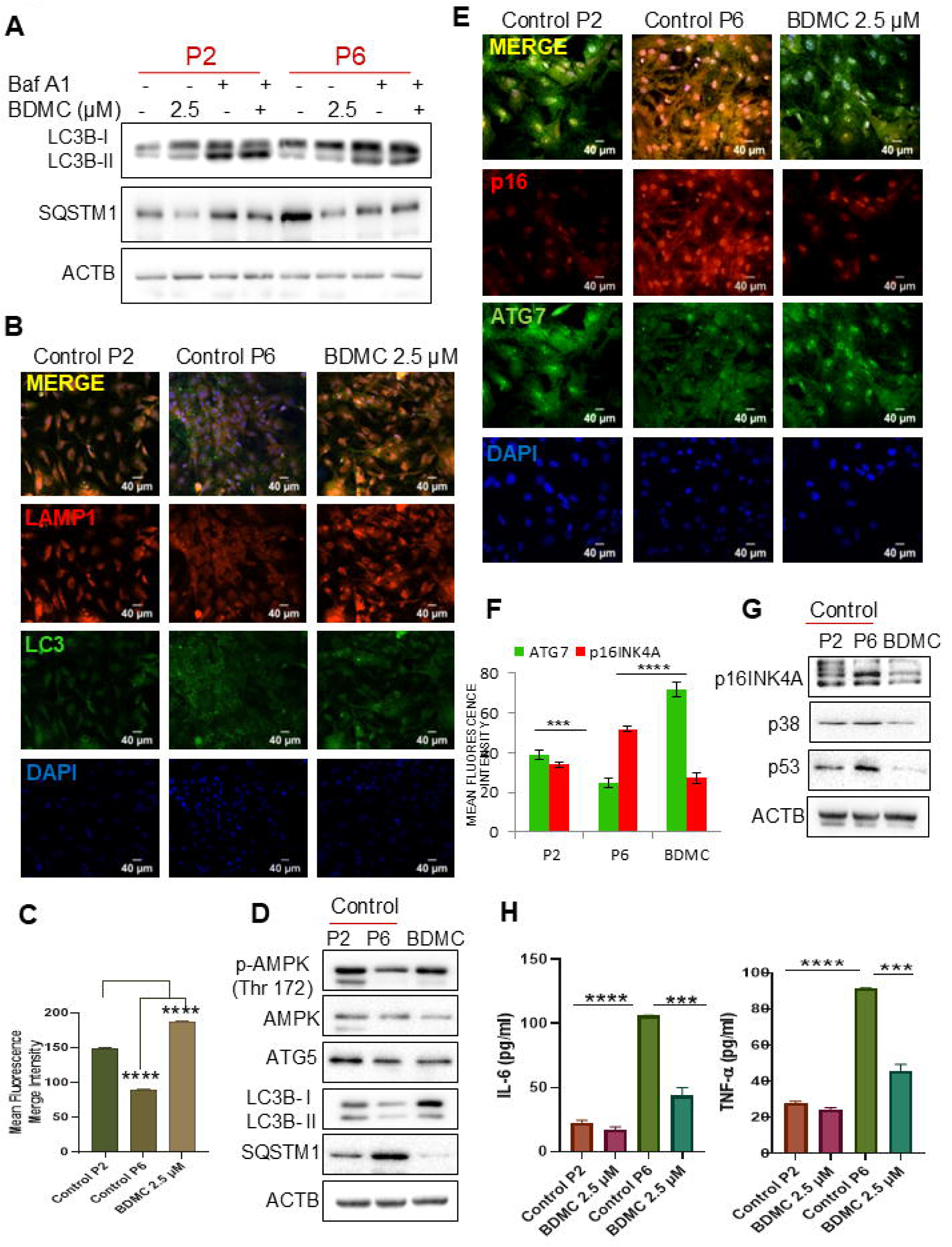
Prolonged treatment of aged astrocytes with BDMC reverses defective autophagy and delays senescence. Astrocytes were pretreated with BDMC through P4 to P6, and comparisons were made with P2 (young) and P6 (aged) astrocytes **(A)** Autophagy flux levels were measured through western blotting in passage 2 and passage 6 astrocytes after treatment with BDMC in presence and absence of Bafilomycin A1 for 3 h. **(B)** Immunoflourescent images depicting LAMP-LC3 colocalisation in young and aged primary astrocytes after prolonged treatment with BDMC.**(C)** Graph representing mean merge fluorescence of 3 (C) Figure. **(D)** Expression levels of autophagy related proteins in young and aged primary astrocytes treated with BDMC represented through western blot. Primary astrocytes at passage 2 and passage 6 and treated with BDMC (p4-p6) were analyzed for the levels of p16INK4A and ATG7 protein. **(E)** Representative confocal fluorescent images depicting levels of p16INK4A and ATG7. Scale bar 40 μm. **(F)** Graph depicting mean fluorescence intensity of 3 (E) Figure. **(G)** Western blot analysis for senescence associated proteins after treatment with BDMC**. (H)** Levels of SASPs measured through ELISA represented by levels of IL-6 and TNF-α. Data were analyzed through one way ANOVA analysis and Bonferroni *post hoc* test. p values ***p < 0.001, **p < 0.01, *p < 0.05

Furthermore, the influence of BDMC on autophagy in aged astrocytes was evaluated through LAMP1-LC3B colocalization. They displayed low levels of LAMP1-LC3B colocalization, and our results showed that senescent astrocytes, when treated with BDMC, exhibited higher immunofluorescence as indicated by higher mean fluorescence intensity (MFI) of LAMP1 and LC3B in treated astrocytes. (Figure 3B-C). These findings signify that with age, dysfunctional autophagic vacuoles marked with LC3B are not able to fuse with lysosomes, causing a reduction in the clearance of autophagic vacuoles loaded with misfolded protein. However, with intervention of BDMC, LAMP1-LC3B colocalization was restored significantly (Figure 3C). We further examined whether senescent astrocytes, which expressed reduced levels of autophagy pathway proteins, could be corrected through BDMC intervention and if it could significantly restore their health. Through western blotting, we found that aged astrocytes with BDMC intervention markedly showed a significant increase in levels of AMPK, an inducer of autophagy, ATG5 involved in an autophagosome formation and LC3B-II, a marker for autophagy and reduced levels of SQSTM1 (Figure 3D and Supplementary Figure S3B).

We also investigated whether dysregulated autophagy was the underlying cause of senescence in aged astrocytes. Through our investigation, we found that senescent astrocytes show a loss in the levels of ATG7 (crucial for the formation of autophagosome) and increased p16INK4A levels, a marker for senescence (Figure 3E). However, BDMC treatment reestablished the levels of ATG7 protein in aged astrocytes and the mean fluorescence intensity for p16INK4A protein was reduced to a significant amount comparable to the levels in young astrocytes (Figure 3E-3F). The effect of BDMC on senescent marker proteins was further confirmed through western blot (Figure3G).The levels of p16, p38 and p53, senescent-associated proteins involved in cell cycle arrest, DNA damage and cellular senescence, were sharply reduced with BDMC intervention compared to their levels in aged astrocytes (Figure 3G and Supplementary Figure S3C).

Next, we checked for the levels of Senescent Associated Secretory Phenotypes (SASPs), which are a collection of factors that promote senescence and have been evidently associated with neuroinflammation in AD pathology ^15^.Our results from ELISA indicated that IL-6 and TNF-α, both prominent SASP factors, were decreased to levels less than half with the use of BDMC when compared to senescent astrocytes. (Figure 3H). Our results hence revealed that the autophagy upregulation by BDMC plays a crucial role in alleviating senescence.

### BDMC restores mitochondria and lysosomal defects caused by senescence

Loss of autophagy with age and in neurodegenerative disorders causes mitochondrial dysfunction and also impairs lysosome-mediated clearance. This is known to lead to the build-up of autophagy by-products, which further exacerbate the disease condition. Several studies have linked loss of autophagy to impairment in lysosomal and mitochondrial function, hence causing senescence ^16^.BDMC was found to increase the levels of mitophagy-associated proteins (Supplementary Figure S4A-B) Through our further study, we were able to find that in senescent astrocytes, the mitochondrial membrane potential was reduced revealed by the low mean fluorescence intensity of Mitotracker red (Figure 4A-B Supplementary Figure S4C) thereby suggesting that with senescence mitochondrial membrane potential also gets affected. Furthermore, prolonged treatment with BDMC was able to reverse this effect and increase the proportion of healthy mitochondria measured by the increased MFI through confocal microscopy (Figure 4A). Next, we analyzed the effect of BDMC on ROS production, which arises because of dysfunctional mitochondria. We assessed higher levels of ROS in senescent astrocytes, and BDMC consequently diminished ROS, the levels of which were measured by DCFH2-DA dye (Figure 4C and Supplementary Figure S4D). These data suggest the positive role played by BDMC in improving overall mitochondrial health.

**Figure 4:**
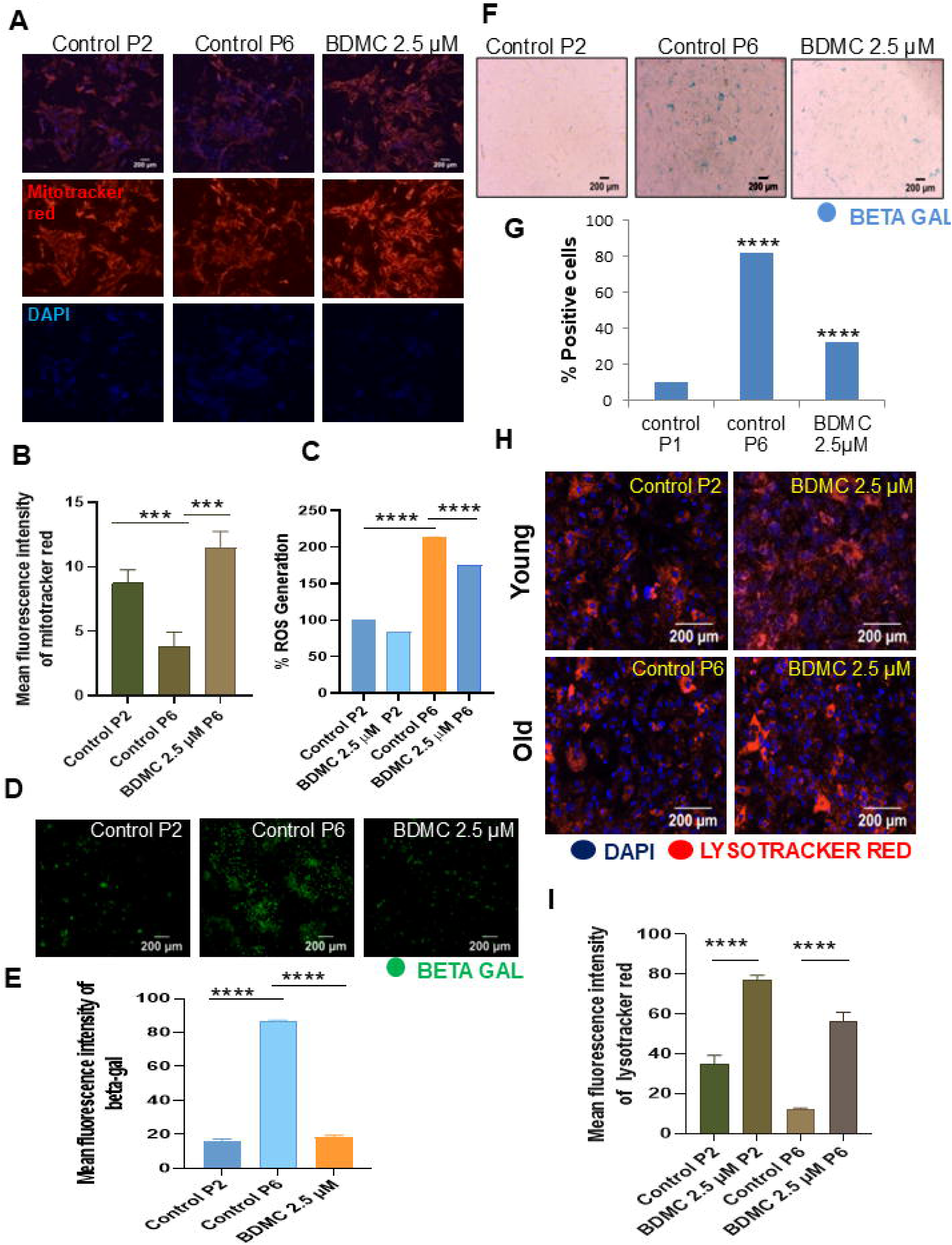
Reduced mitochondrial and lysosomal function of aged astrocytes is restored with prolonged treatment with BDMC **(A)** Mitochondrial membrane potential was measured through levels of Mitotracker Red in young, aged, and BDMC treated astrocytes. **(B)** Graph depicting mean fluorescence intensity of 4 (A) **(C)** ROS quantification in young and treated aged astrocytes measured through levels of DCFH-2DA. **(D)** Representative Immunofluorescence images of assay beta-gal staining of primary astrocytes treated similarly as in Fig 3. **(E)** Graph depicting mean fluorescence intensity of 4 (D) **(F)** Representative images for SA-β-Gal staining in young aged and BDMC treated primary astrocytes**. (G)** Graph showing number of SA-β-Gal positive cells. **(H)** Representative confocal images of young and aged astrocytes depicting levels of lysotracker red after treatment with BDMC and lysosome inhibitor bafilomycin A1**. (I)** The representative graph showing mean fluorescence intensity for 4(G). Data were analyzed through one-way ANOVA analysis and Bonferroni *post hoc* test. p values ***p < 0.001, **p < 0.01, *p < 0.05

With disrupted autophagy, lysosomes also accumulate and are not able to fuse with the autophagosome during AD^2^. Reduced lysosomal activity is a feature of defective autophagy and neurodegenerative disorders occurring with age. For these reasons, we next examined the effect of BDMC on beta-galactosidase activity, which is a well-established marker for senescence and unhealthy lysosomal activity (Figure 4D-E). Beta-galactosidase-positive cells were notably increased in senescent astrocytes when compared to young astrocytes (Figure 4D-G). Correspondingly, with BDMC, the percentage of SA-b-Gal-positive cells decreased drastically, measured by MFI and the number of SA-b-Gal-positive cells (Figure 4E and 4G), suggesting the positive role of BDMC in alleviating senescence-associated beta-galactosidase activity of lysosomes. Furthermore, our results indicated that BDMC enhanced the acidity of lysosomes, which was otherwise reduced in aged astrocytes and is associated with AD pathology. (Figure 4H). The mean fluorescence intensity of mitotracker red used to mark acidic lysosomes showed a substantial increase in BDMC-treated senescent astrocytes (Figure 4H-I).The above results indicate that the positive effect of BDMC on mitochondria and lysosome health is relevant to its role in delaying senescence and correcting defects in autophagy.

### Genetic impairment of AMPK-mediated autophagy reverses the effect of BDMC on senescence

Through our above studies, we confirmed that BDMC is able to alleviate senescence and also enhance autophagy in senescent primary astrocytes. To further confirm the relationship between the pathways of autophagy and senescence and establish the underlying mechanism involved in the role of BDMC in augmenting autophagy and reducing senescence, AMPK, a key regulator of autophagy was knocked down from both young and senescent astrocytes (Supplementary Figure S5A-B) For this study we transfected astrocytes with *siRNA* for AMPK and checked its effect on senescence inhibition activity of BDMC. We found that knockdown of AMPK reversed the autophagy induction in aged astrocytes by BDMC. Through western blot assay, we found that the levels of LC3B-II were reduced and SQSTM1 were increased in BDMC-treated senescent astrocytes where AMPK was knocked down (Figure 5A and supplementary Figure S5C). AMPK knockdown also reversed the anti-senescence activity of BDMC, and levels of p16 and p38 increased in treated senescent astrocytes. (Figure 5A and supplementary Figure S5C).

**Figure 5:**
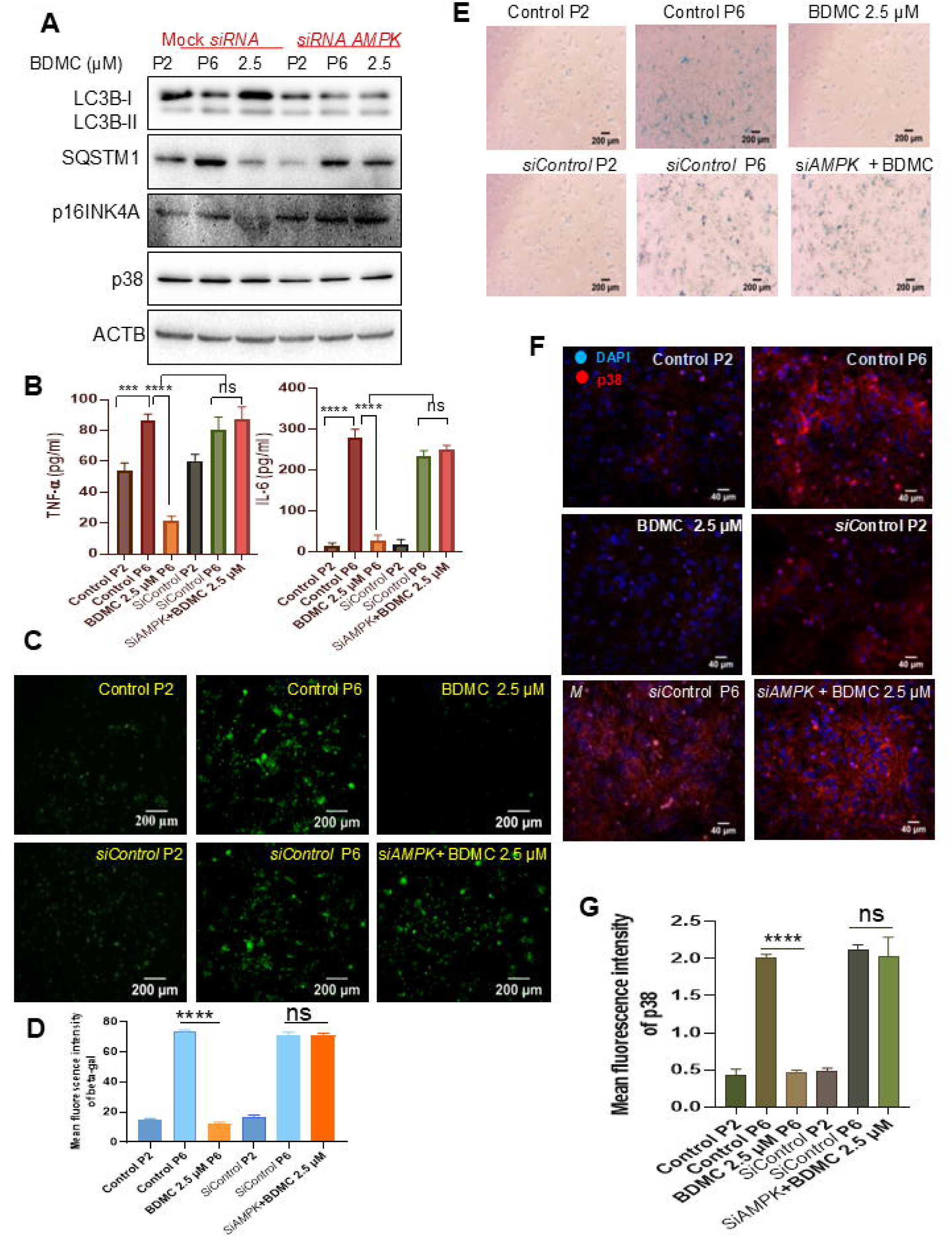
BDMC delays senescence in aged astrocytes in AMPK dependent manner. Expression of AMPK protein was knocked down from primary astrocytes through siRNA transfection of cells. (A) Immunoblots of autophagy and senescence-associated proteins after the given treatment. (B) Expression levels of SASP TNF-[ and IL-6 measured through ELISA (C) Representative confocal images of SA-β-GALstaining. Scale bar 200 μm. (D) Graph representing SA-β-GAL positive astrocytes after treatment with BDMC and reduced expression of AMPK. (E) Representative images of SA-β-GAL staining in AMPK knockdown astrocytes after treatment with BDMC. (F) Immunostaining analysis of AMPK silenced aged astrocytes for expression levels of senescence associated protein p38 levels. Scale bar 40 μm. (G) Graphical representation of the mean fluorescent intensity of Figure 5 (F) Data analyzed through one way ANOVA and Bonferroni *post hoc* tests. p values ***p < 0.001, **p < 0.01, *p < 0.05

Furthermore, the influence of AMPK on senescence-associated beta-gal was assessed. We found a similar trend that with reduced expression of AMPK from astrocytes, levels of SASP cytokines IL-6 &TNF-α also increased, and BDMC-treated cells showed signs of senescence (Figure 5B). The effect of BDMC on the senescence of aged astrocytes was hugely blunted because of AMPK knockdown, as indicated by the increased number of SA-β Gal positive cells in BDMC-treated astrocytes (Figure 5C-5E). Interestingly, reduced AMPK expression in cells also led to an increase in levels of protein p38 (Figure 5F-G). This was confirmed by Immunofluorescence of p38 in treated cells, indicated by mean fluorescence intensity (MFI) (Figure 5F-G). These findings indicate that autophagy mediated through AMPK is the pathway that confers BDMC the ability to cause depletion of senescence-associated isoforms in aged astrocytes.

### BDMC cleared Aβ_42_ in primary astrocytes by inducing autophagy through the AMPK pathway

Defective autophagy contributes to the accumulation of amyloid beta in AD. To evaluate the effect of BDMC on amyloid beta clearance, we preincubated astrocytes with BDMC for 12h, followed by incubation with Aβ_42_ for 12 h. We found that the astrocytes treated with BDMC displayed a significant clearance of Aβ_42_ (Figure 6A).

**Figure 6:**
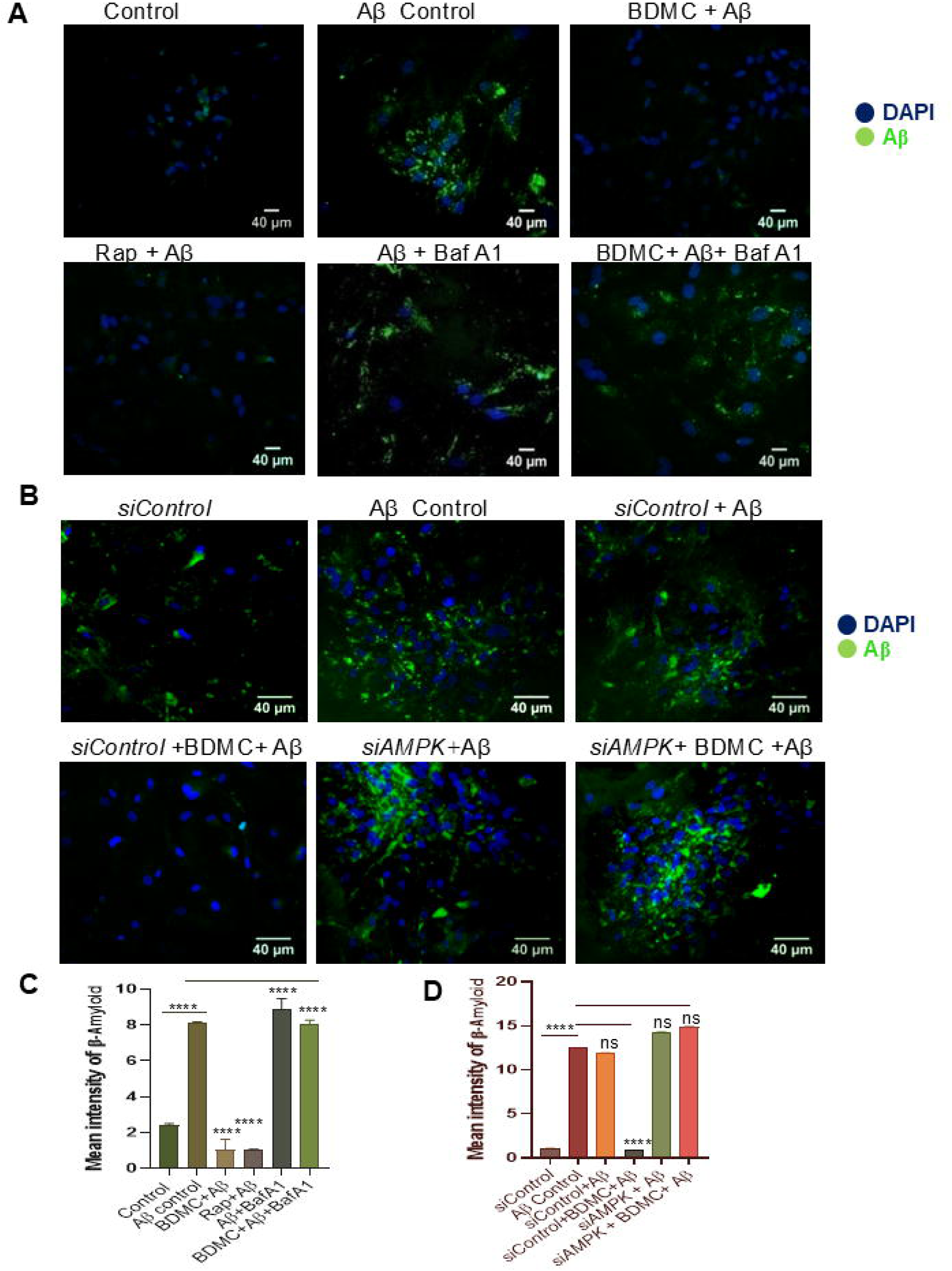
BDMC promotes Aβ_42_ clearance through AMPK-mediated autophagy. Astrocytes were treated for 12 h with 2 μg/ml of fluorescently tagged Aβ (hilyte 488) after BDMC treatment to cells at 2.5 μM conc. **(A)** Representative confocal images of Aβ_42_ deposits and effect of BDMC on the Aβ_42_ clearance in primary astrocytes. To reduce the expression of AMPK from cells, primary astrocytes were transfected and treated similarly as in Figure 6(A).**(B)** Immunoflourescent images showing expression levels of Aβ_42_ in primary astrocytes with reduced expression of AMPK **(C)**Mean fluorescent intensity of Figure 6(A).**(D)** Graph representing Mean fluorescent intensity of Figure 6(B). Data were analyzed through one-way ANOVA and Bonferroni *post hoc* test was applied for statistical comparison. p values ***p < 0.001, **p < 0.01, *p < 0.05

The confocal microscopy data revealed that astrocytes treated with BDMC and rapamycin (a known autophagy inducer) showed clearance of Aβ_42_ to a greater degree (Figure 6A and 6C). The reversal in this effect was seen when autophagy inhibitor bafilomycin A1 was added for 3 h, indicating the role of autophagy in amyloid beta clearance (Figure 6A).

To further validate our findings, we genetically knocked down AMPK in astrocytes to inhibit autophagy. In the presence of *siAMPK*, BDMC could not clear amyloid beta from cells, suggesting that autophagy mediated through the AMPK pathway is responsible for this potential of BDMC to clear amyloid beta (Figure 6B and 6D) and that an autophagy defect hampers amyloid clearance by BDMC.

### BDMC improved spatial memory, exploratory behavior and neuromuscular coordination in 3xTg-AD mice

After confirming the effect of BDMC on autophagy-mediated inhibition of senescence in the primary astrocytes, we attempted to validate our findings in the transgenic mouse model of AD (3xTg-AD). However, before going for the disease model, we tested the bioavailability of BDMC across the BBB. The pharmacokinetic study of BDMC was performed in C57BL/6J mice after oral administration of BDMC at 50 mg/kg. The mean plasma concentration versus time profiles and mean brain concentration (ng/ml) of BDMC are represented in Figure 7A and 7B, respectively. BDMC could reach the highest plasma concentration of 113 ng/ml within a time frame of 15 minutes, and the mean brain concentration after BDMC was found to be 20 ng/g (SupplementaryTable 1).Overall, the data suggested that the BDMC had satisfactory pharmacokinetic behavior and was sufficiently available across the BBB for further studies. Henceforth, we treated 6-month-old 3xTg-AD mice for the duration of two months and analyzed different behavioral parameters through various tests. In the radial arm maze test, the untreated 3xTg-AD mice displayed a significant reduction in spatial memory in comparison to wild-type controls, as the time taken to reach the baited arm was longer(Figure 7C-D). However, BDMC-treated groups at 50 and 100 mg/kg dose took significantly less time with reduced distance traveled to find their bait with the help of a visual cue. BDMC-treated groups spent more time in the baited arm, apart from fewer entries into non-baited arms, in comparison to untreated 3xTg-AD mice, indicating improvement in spatial memory (Figure 7C-D). In the open field test, untreated 3xTg-AD explored less as their time mobile and maximum speed was less in comparison to WT control; their center zone time was more rather than spending time in corners (Figure 7E-F). Furthermore, the mice treated with BDMC displayed significant improvement in locomotor and exploratory activity as well (Figure 7E-F). In the rota rod test, 3xTg-AD mice showed poor neuromuscular coordination as their latency to fall from a rotating rod was shorter, whereas BDMC-treated groups had a significant increase in latency to fall from a rotating rod (Figure 7G), displaying better neuronal function and improved motor coordination.

**Figure 7:**
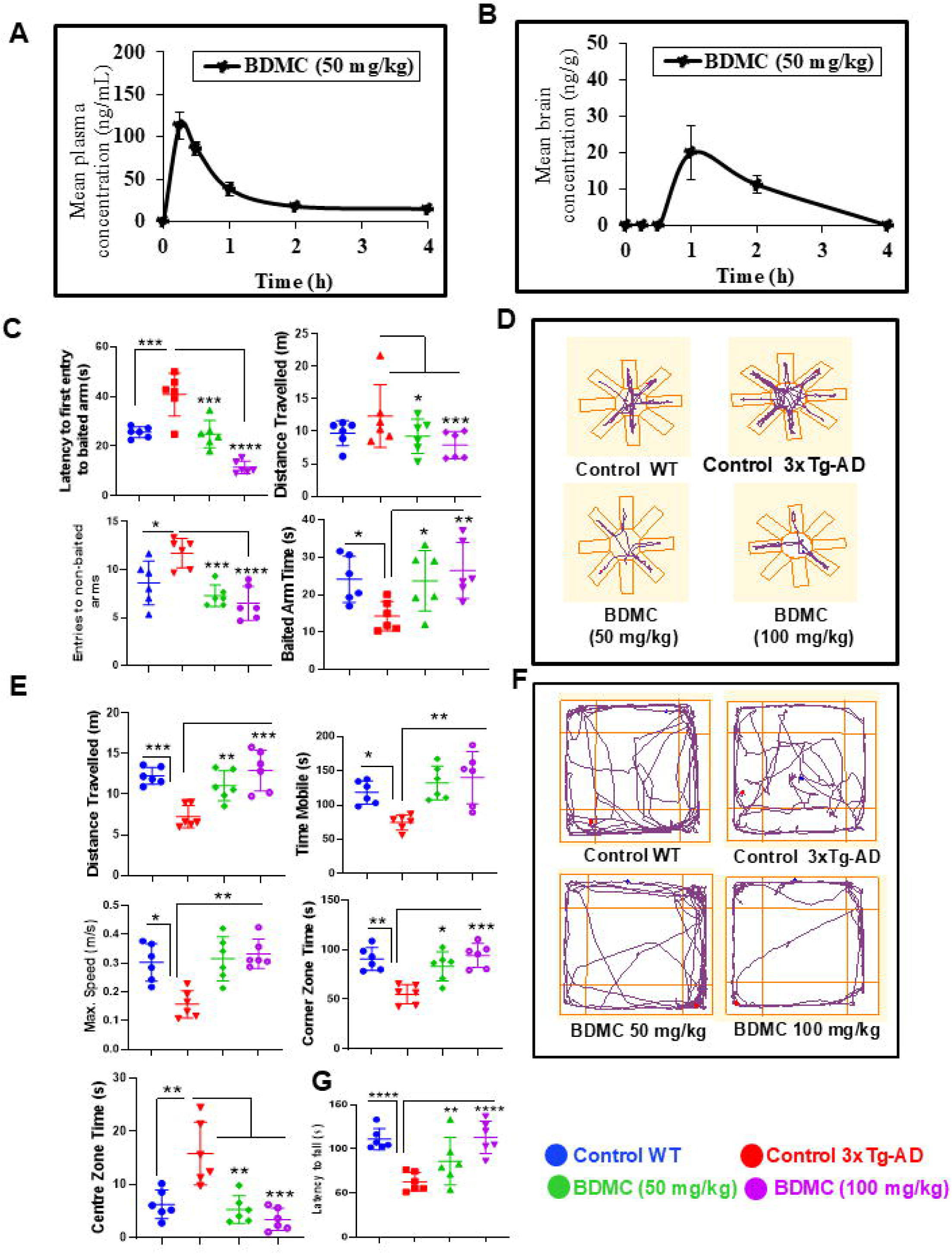
BDMC alleviates AD pathophysiology in 3xTg-AD mice. Six months old 3xTg-AD mice were administered BDMC orally at 50 mg/Kg and 100 mg/Kg for 2 months. **(A)** Pharmacokinetic analysis: Mean plasma concentration of BDMC in C57 BL6/J mice after dosing of 50 mg/kg. **(B)** Brain concentration of BDMC.BDMC reverses cognitive impairment in 3xTg-AD mice. To assess the effect of BDMC on AD pathophysiology, Open Field Test (OFT), Radial Arm maze and Rotarod test were performed. **(C)** Graphs representing latency to first entry to baited arms, distance travelled, entries to non-baited arms and baited arm post five days of training. **(D)** Track plots of representative RAM analyzed by ANYmaze software. **(E)** Total distance traveled, time mobile, maximum speed frequency of entries in center and latency of center entry in Open field test. **(F)** Representative track plots of OFT. **(G)** Latency to fall time of mice assessed by Rota rod test. All data is represented as mean (±SEM). Two-way ANOVA was used to compare group treated with BDMC effect between WT and 3xTg-AD mice, followed by a Bonferroni test as a post hoc test for statistical analysis. p values ***p < 0.001, **p < 0.01, *p < 0.05. (n=7)

### BDMC reduced the Aβ_42_ plaques, the number of reactive astrocytes and neuroinflammation in 3xTg-AD mice

Behavioral changes in 3xTg-AD mice after treatment with BDMC led us to investigate the underlying cause for those alterations. Therefore, through immunohistochemistry (IHC), we checked the levels of Aβ_42_ in the hippocampus after two months of treatment with BDMC. We found a significant decline in the levels of Aβ_42_ in the hippocampus at both doses, 50 mg/kg and 100 mg/kg of BDMC, respectively, compared to the control group, as indicated by brown-colored plaques (Figure 8A). The average number of Aβ_42_ plaques was significantly reduced in BDMC-treated groups as compared to the 3xTg-AD control group (Figure 8C). Aβ_42_ is strongly related to activation of astrocytes, we therefore checked if reduced levels of Aβ_42_ had any impact on the reactive phenotype of astrocytes. The expression of GFAP was reduced with BDMC treatment, as indicated by reduced brown pigmented depositions (Figure 8B), and the number of GFAP+ astrocytes in the field was also reduced significantly at both doses. (Figure 8D), Further, the reduction of Aβ_42_ levels in the brain also led to reduced levels of proinflammatory cytokines TNFα, IL-6 and IL-1β in the cortex region of the brain of 3xTg-AD mice treated with BDMC in comparison to untreated control mice, as observed through ELISA (Figure 8E), which clearly established the strong effect of BDMC against neuroinflammation and astrogliosis.

**Figure 8:**
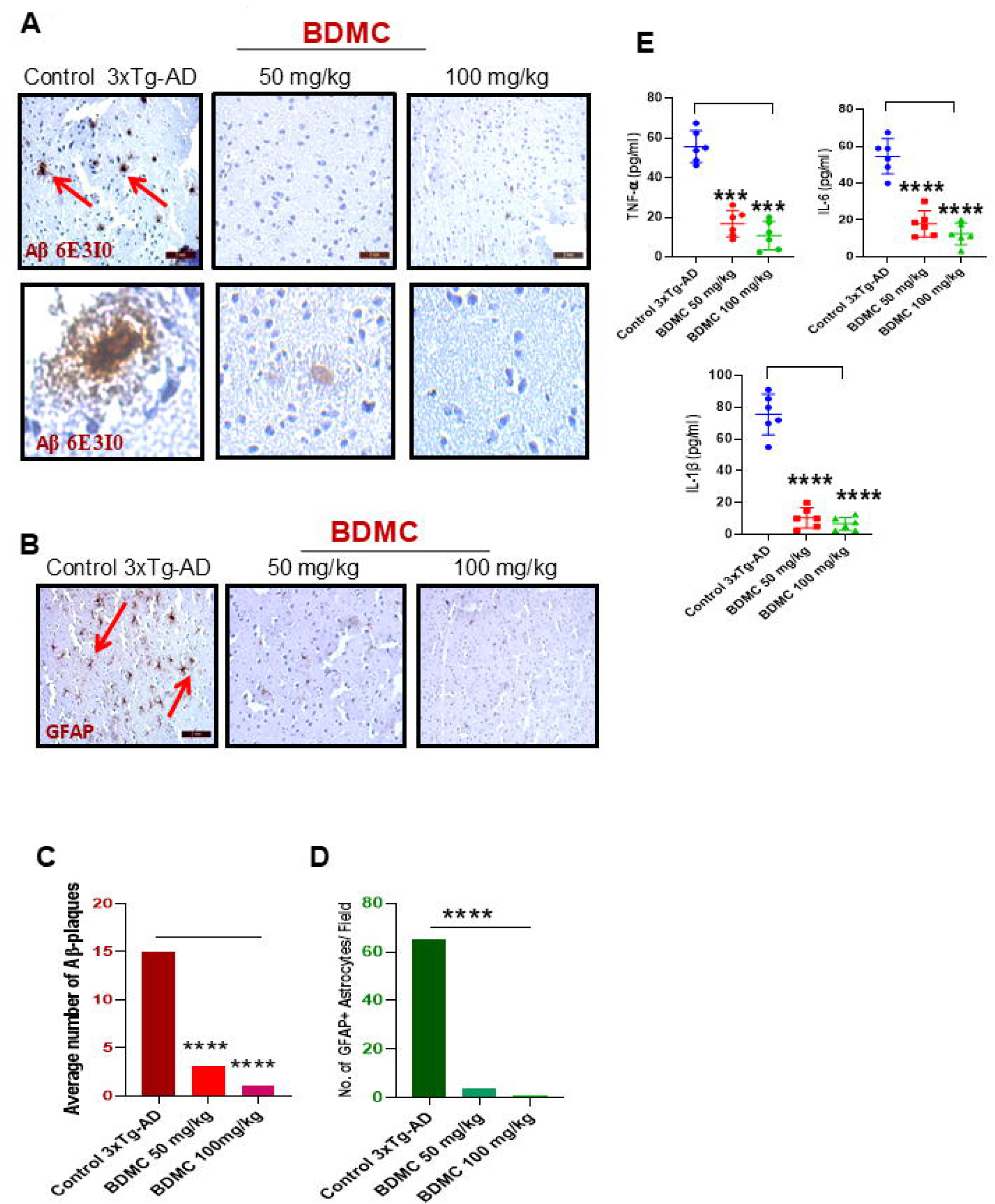
BDMC ameliorates levels of AD markers in Hippocampal region of 3xTg-ADmice brain. 3xTg-AD mice were treated as in Figure 7. **(A)** Representative IHC images of Aβ deposits (brown) in hippocampal region of 3xTg-AD mice (6 month old) treated with BDMC at 50 mg/kg and 100 mg/kg for two months**. (B)** IHC images representing levels of astrocytic marker GFAP in 3xTg-AD brain. **(C)** Quantification of Aβ plaques**. (D)** Levels were quantified by measuring the number of GFAP positive astrocytes in hippocampi of 3xTg-AD mice**. (E)** Relative levels of TNF-α, IL-6 and IL-1β in brain of 3xTg-AD mice measured by ELISA. Statistical analyses were done using one-way ANOVA and Bonferroni test for multiple comparisons. p values ****p <0.0001,***p < 0.001. (n=7)

### BDMC restored autophagy and alleviated senescence in 3xTg-AD mice

To further substantiate our findings from *in vitro* data that BDMC alleviates senescence and mitigates defects of autophagy, we next analyzed the effect of BDMC on key proteins involved in autophagy and senescence in the 3xTg-AD mice brain hippocampi. Consistent with our previous data, BDMC increased the levels of key autophagy pathway proteins pAMPK, ATG5, ATG7, and BECN1 analyzed through western blotting (Figure 9A-B). The expression for SQSTM1 was reduced in both 50mg/kg and 100mg/kg groups (Figure 9A-B). Similarly, p16, p38 and p53 depicted increased levels in control 3xTg-AD mice and BDMC at 50 and 100 mg/kg significantly minimized their expression. (Figure 9C-D) Thus, loss of autophagy drives senescence in 3xTg-AD mice, and BDMC intervention was able to restore the autophagy and delay senescence.

**Figure 9:**
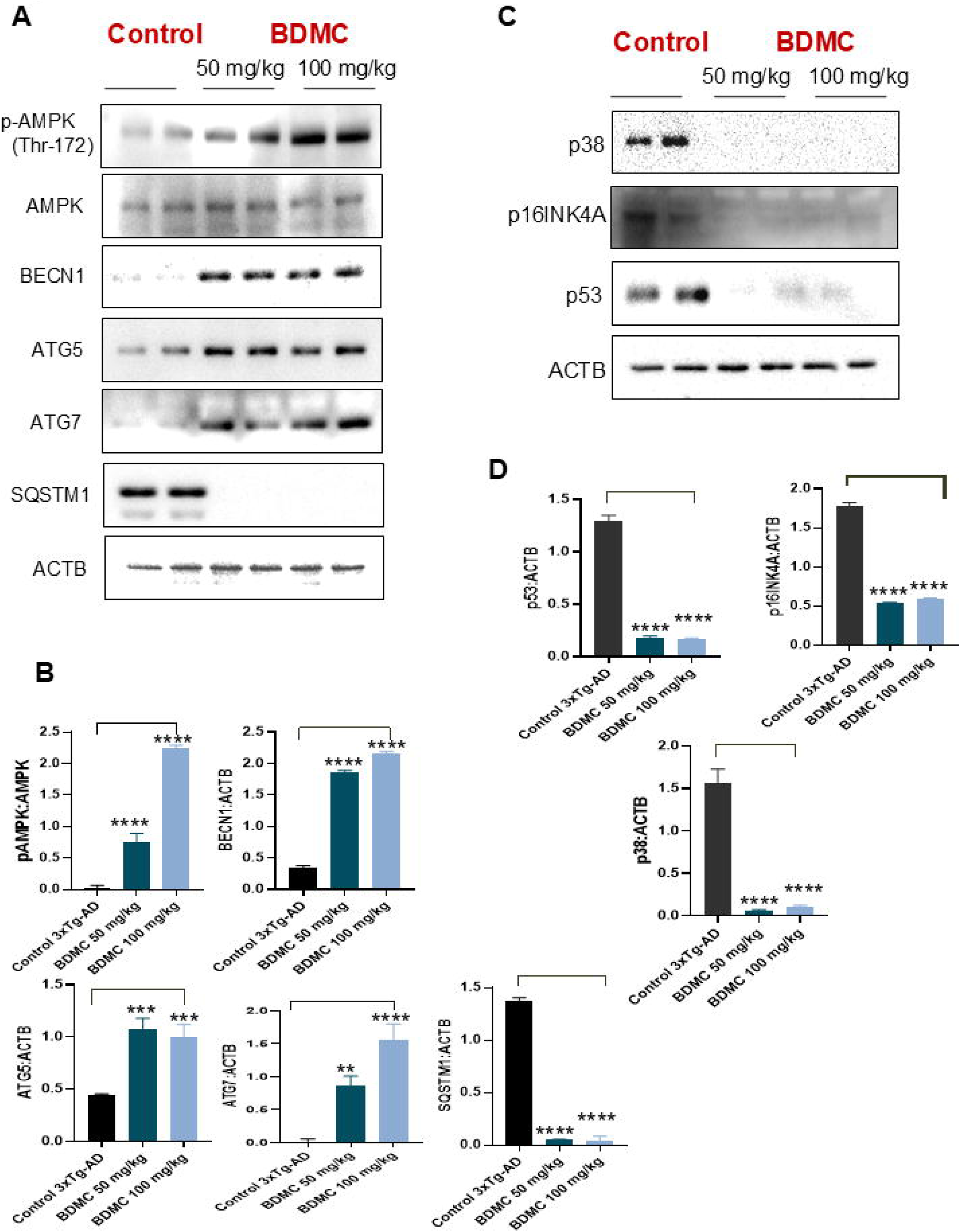
BDMC effectively upregulated autophagy and delayed senescence cascade in brain of AD mice. **(A)**Representative western blot images of autophagy related proteins. **(B)** Quantification of the protein levels normalized with ACTB. **(C)** Immunoblots depicting expression levels of senescence related proteins p38, p16INK4A and p53. **(D)** Quantification of the protein levels expression normalized with ACTB. Statistical significance was analyzed using one way ANOVA and Bonferroni test for multiple comparisons. p values ****p <0.0001,***p < 0.001. (n=7).

## Discussion

Aging is a key driver for the development of Alzheimer disease^17^. It is a complex biological process that affects various cellular mechanisms, including autophagy, which is essential for maintaining cellular homeostasis^18^. A gradual decline in the expression of autophagic proteins is strongly related to the pronounced increase in senescence proteins and reduced functional capabilities of brain cells ^19^. These age-related changes in the brain cells contribute significantly to reduced clearance of amyloid beta from the brain and compromised functionality^20^. In this study, we attempted to capture these changes in vitro in the aging primary astrocytes and in vivo in the 3xTg-ADmice, a transgenic mouse model for AD. We demonstrated that the aging of primary astrocytes is accompanied by a significant reduction in autophagy, marked by decreased levels of key autophagic proteins such as ATG5, ATG7, BECN1, LC3BII, and increased expression of SQSTM1. This reduction in autophagy flux correlates with the rise in cellular senescence, as evidenced by elevated levels of various senescence markers, including p16INK4A, p38, and p53, alongside increased β-galactosidase activity. These findings suggest that the ageing process leads to impaired autophagy in astrocytes, contributing to a cellular environment conducive to senescence. Moreover, our results highlight a potential mechanism linking the decline in autophagy to the increased incidence of senescence in aged cells, a hallmark feature of neurodegenerative diseases like Alzheimer’s disease (AD)^21^.

Given the central role of autophagy in cellular maintenance and the detrimental effects of its dysfunction in aging and neurodegeneration, we sought to investigate potential therapeutic strategies to restore autophagy and mitigate senescence in aged astrocytes. We identified bisdemethoxycurcumin curcumin (BDMC), a curcuminoid derived from *Curcuma longa*, as a potent inducer of autophagy. Our results showed that BDMC treatment significantly increased autophagy in aged astrocytes, as evidenced by elevated levels of autophagy markers such as LC3B-II and a decrease in SQSTM1. Additionally, BDMC treatment upregulated key autophagy-related proteins such as pAMPK, pULK1, FIP200, BECN1, and ATG5, while reducing the levels of mTOR, a negative regulator of autophagy. Importantly, BDMC treatment did not impair cell viability, even after prolonged exposure, suggesting that it is a safe and effective compound for restoring autophagy in astrocytes. These findings support BDMC as a promising candidate for targeting autophagy dysfunction in age-related neurodegenerative diseases.

We further asked if the induction of autophagy by BDMC can help alleviate senescence in aged astrocytes. Our data demonstrated that BDMC treatment not only restored autophagy flux but also reduced the accumulation of senescence markers. Specifically, BDMC reduced the expression of p16, p38, and p53, the key markers associated with cellular senescence, and lowered the levels of SASP factors,including IL-6 and TNF-α, which are linked to neuroinflammation. The release of SASPs by senescent cells acts as a driving force for neurodegenerative damage, which further contributes to the pathology of diseases like AD^22^. Therefore, the ability of BDMC to enhance autophagy may play a key role in mitigating senescence and associated neuroinflammation, which are prominent features of neurodegenerative diseases like AD^23^.

The impact of BDMC on mitochondrial and lysosomal function in aged astrocytes further supports its potential as a therapeutic agent. Mitochondrial dysfunction and lysosomal impairment are commonly observed in aging and neurodegenerative diseases, and both are closely linked to defective autophagy^8^. Our data demonstrated that BDMC restored mitochondrial membrane potential and reduced ROS production, both of which were impaired in senescent astrocytes. Additionally, BDMC treatment improved lysosomal activity, as evidenced by reduced β-galactosidase activity and restored lysosomal acidity in aged astrocytes. These findings indicate that BDMC not only enhances autophagy but also contributes to the overall health of mitochondria and lysosomes, which are critical for cellular homeostasis.

To establish the underlying mechanism of BDMC’s effects, we investigated the role of AMPK, a key regulator of autophagy, in mediating the compound’s effects on autophagy and senescence. This pathway is the key regulator of energy metabolism in the cells and gets activated physiologically when ATP levels are reduced due to its conversion to ADP and AMP^24^. However, this pathway can be activated pharmacologically and can help effectively clear amyloid beta from the brain of the transgenic AD mouse model 5XFAD, as we have demonstrated earlier ^25^.

Our results showed that the knockdown of AMPK in astrocytes reversed the beneficial effects of BDMC on autophagy and senescence. Specifically, AMPK knockdown reduced the levels of LC3B-II and increased the levels of SQSTM1, while also reversing the reduction in senescence markers such as p16 and p38. These results suggest that the autophagy-inducing effects of BDMC are mediated through AMPK signaling and that AMPK plays a crucial role in BDMC’s ability to mitigate senescence in astrocytes.

The therapeutic potential of BDMC was further validated in a transgenic mouse model of Alzheimer’s disease (3xTg-AD mice). However, before validating the efficacy of BDMC in 3xTg-AD mice, we confirmed its bioavailability in both plasma and the brain. We found a significant quantity of BDMC in both plasma and the brain, unlike other curcuminoids like curcumin and demethoxycurcumin, for which poor bioavailability is the biggest issue ^26^. Further, our data showed that chronic BDMC treatment of 3xTg-AD mice for two months led to significant improvement in spatial memory, exploratory behavior, and neuromuscular coordination in these mice. The better performance of BDMC-treated mice in these tests clearly indicated improvement in brain health. Furthermore, these behavioral improvements were accompanied by a significant reduction in amyloid beta (Aβ) plaques in the hippocampi of these mice and a reduction in the reactive phenotype of astrocytes.

The improvement in brain health was also accompanied by reduced levels of proinflammatory cytokines TNF-α, IL-6, and IL-1β in the brain cortex, indicating decreased neuroinflammation in BDMC-treated mice. Further investigation revealed that, similar to its pharmacological effects in primary astrocytes, BDMC treatment led to the upregulation of autophagic markers and downregulation of senescence markers in the hippocampi of 3xTg-AD mice. These findings indicate that BDMC not only restores autophagy and reduces senescence in vitro but also improves cognitive function and reduces neuroinflammation in vivo, suggesting its potential as a therapeutic agent for AD.

In conclusion, our study provides compelling evidence that BDMC, a curcuminoid derived from *Curcuma longa*, can effectively restore autophagic function, reduce senescence, and alleviate the pathological features of Alzheimer’s disease in both primary astrocytes and in the 3xTg-AD mouse model. The therapeutic effects of BDMC appear to be mediated through the activation of the AMPK pathway, highlighting the potential of BDMC as a novel, non-toxic agent for combating aging-related neurodegenerative diseases. Further studies are warranted to investigate the long-term efficacy and safety of BDMC in clinical settings, but our findings lay the basis for the development of BDMC-based therapies for age-related neurodegeneration.

## Acknowledgments

We are thankful to the Council of Scientific and Industrial Research (CSIR), India, for providing the research fellowship to Ms. Parul Khajuria.

## Ethics Statement

This study used mice. All the necessary approval for the use of animals was taken by the authors from the animals’ ethics committee of CSIR-Indian Institute of Integrative Medicine, Jammu. The authors also declare that all the data shown in this manuscript are true to the best of our knowledge.

## Authors’ contributions

PK and DK did most of the in vitro experiments, along with some in vivo experiments. DK and LS performed ELISA experiments. KS performed animal behavior studies. Razia B and SBB isolated and characterized the compound BDMC. DM and UN did pharmacokinetics studies. PR helped in the genotyping of 3x-Tg mice. ZA helped in designing the study and provided institutional support. PK and AK designed the study, analysed the data, and wrote the manuscript.

## Abbreviations

Aβ: amyloidbeta
AD: Alzheimer disease
AMPK: 5’ adenine monophosphate-activated protein kinase
ATG: autophagy related
BECN1: beclin 1, autophagy related
DAPI: 4,6-diaminido-2-phenylindole
GFAP: glial fibrillary acidic protein
IBA1: ionized calcium-binding adapter molecule 1
BC-i: sobavachalcone
IHC: immunohistochemistry
LAMP-1: lysosomal-associated membrane protein 1
LC3: light chain 3
MTOR: mechanistic target of rapamycin kinase
NeuN: neuronal nuclear protein
NFT: neurofibrillary tangles
OFT: open field test
RAM: Radial Arm Maze
SQSTM1: Sequestosome 1
ULK-1: unc-51-like kinase 1

