## Supplementary Information Figures for "Bisdemethoxycurcumin mitigates Alzheimer disease pathology through autophagy-mediated reduction of senescence and amyloid beta"

### Bis-demethoxy curcumin mitigates AD pathology through autophagy-mediated reduction in senescence in 3XTG mice

#### Supplementary Files

**Fig S1 A**

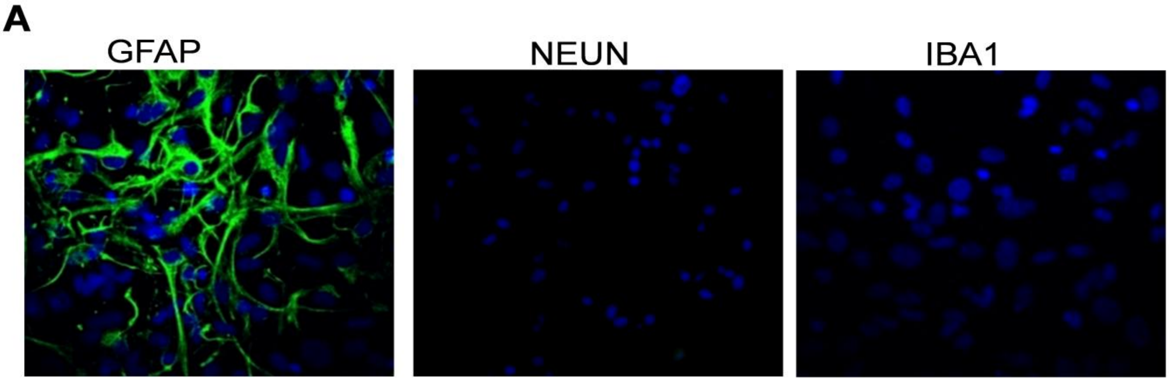


**Figure S1A** Cells were isolated from brain of 3-4 days old C57BL/6J mice pups and analysed for the purification of culture. (A) Immunofluorescence images depicting brain cells where GFAP (green) marker stains astrocytes IBA1 marker stains microglia and NEUN stains neurons and nucleus stained by DAPI (blue). Primary cells isolated from brains of mice pups were tested for expression of all these markers and analysed for purity of astrocytes.

Operator:NPMC Timebase:HPLC Sequence:Caffeine07052024 Page 1-1
11-10-2024 10:54 AM

#### 12 BDMC

| | | | |
|---|---|---|---|
| *Sample Name:* | **BDMC** | *Injection Volume:* | **10.0** |
| *Vial Number:* | **RA6** | *Channel:* | **UV_VIS_3** |
| *Sample Type:* | **unknown** | *Wavelength:* | **254.0** |
| *Control Program:* | **30-80** | *Bandwidth:* | **4** |
| *Quantif. Method:* | **Caffeine07052024** | *Dilution Factor:* | **1.0000** |
| *Recording Time:* | **11-10-2024 10:03** | *Sample Weight:* | **1.0000** |
| *Run Time (min):* | **45.00** | *Sample Amount:* | **1.0000** |

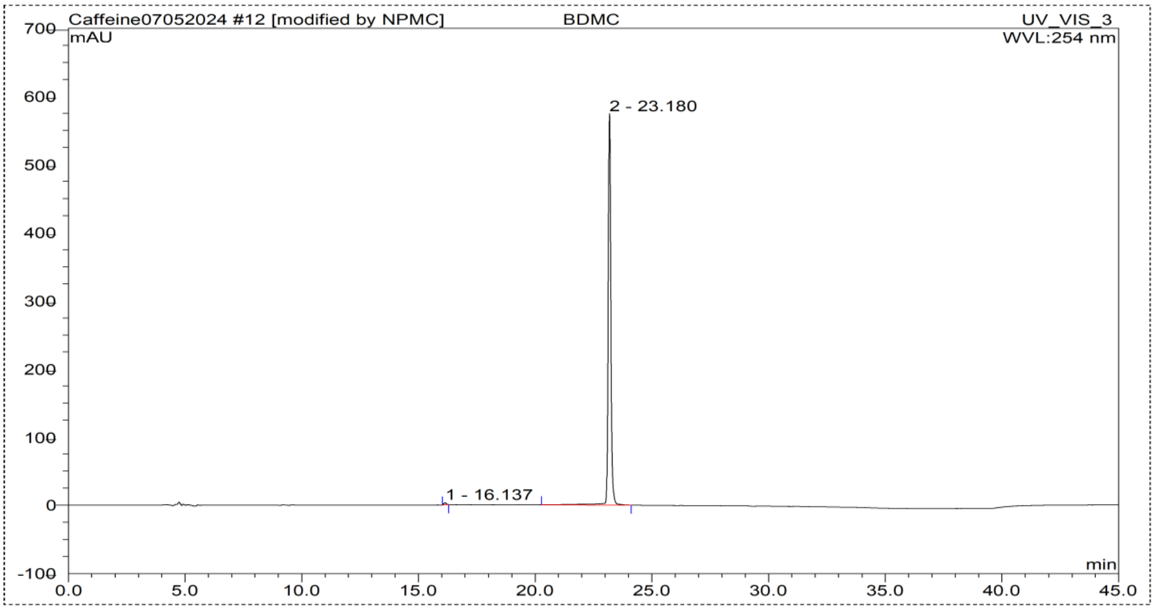


| No. | Ret.Time min | Peak Name | Height mAU | Area mAU*min | Rel.Area % | Amount | Type |
|---|---|---|---|---|---|---|---|
| 1 | 16.14 | n.a. | 3.155 | 0.380 | 0.49 | n.a. | BMB* |
| 2 | 23.18 | n.a. | 573.833 | 76.366 | 99.51 | n.a. | BMB |
| **Total:** | | | 576.988 | 76.746 | 100.00 | 0.000 | |

DEFAULT/Integration

Chromeleon (c) Dionex 1996-2006
Version 6.80 SR15 Build 4656 (243203)

**Figure S1B** Characterisation of BDMC; purity of isolated BDMC (99%), its HPLC and NMR.

**Fig S1C**

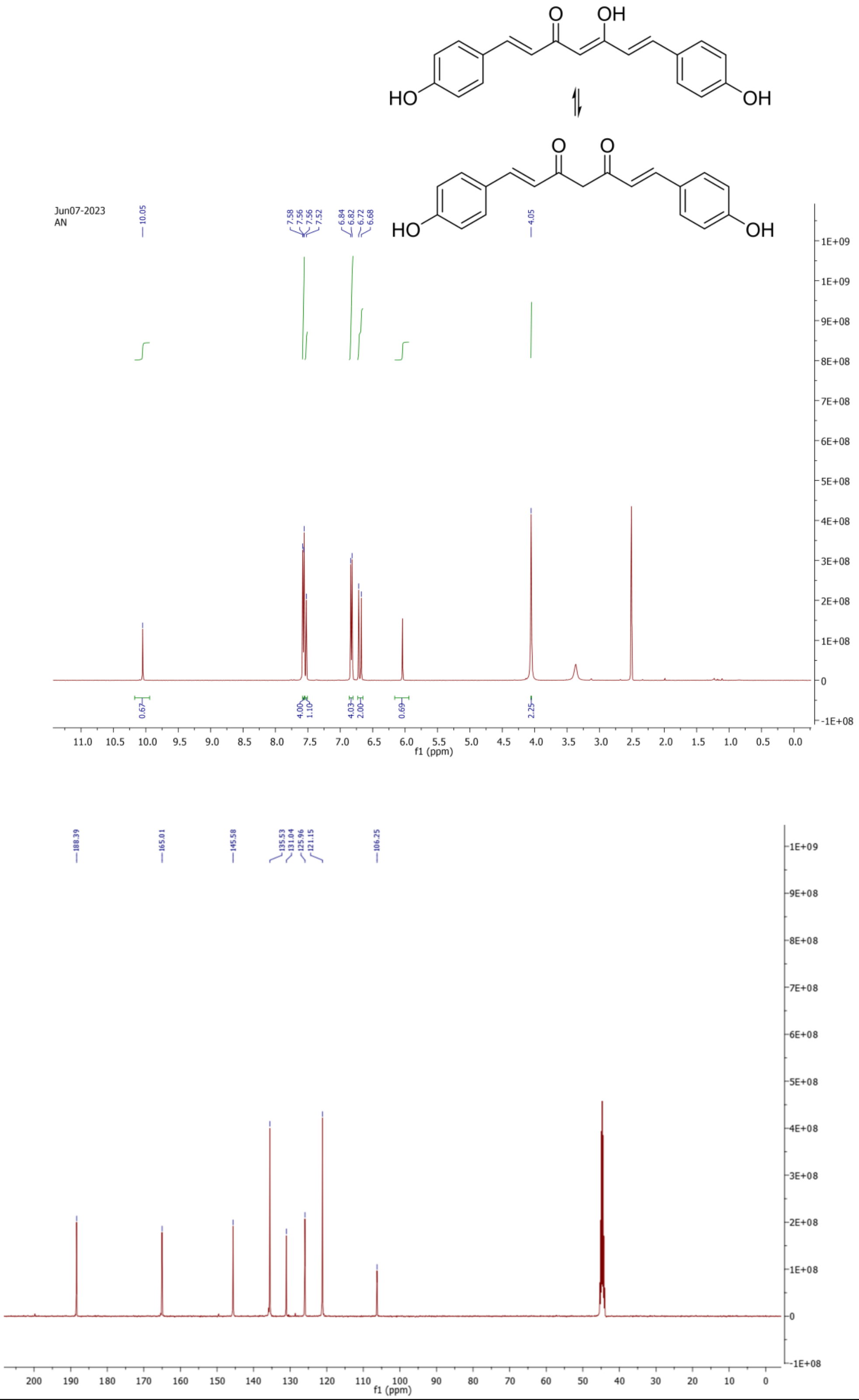
O
OH
HO
OH
O
O
HO
OH
Jun07-2023
AN
10.05
7.58
7.56
7.56
7.52
6.84
6.82
6.72
6.68
4.05
0.67
4.00
1.10
4.03
2.00
0.69
2.25
f1 (ppm)
188.39
165.01
145.58
135.53
131.04
125.96
121.15
106.25
f1 (ppm)

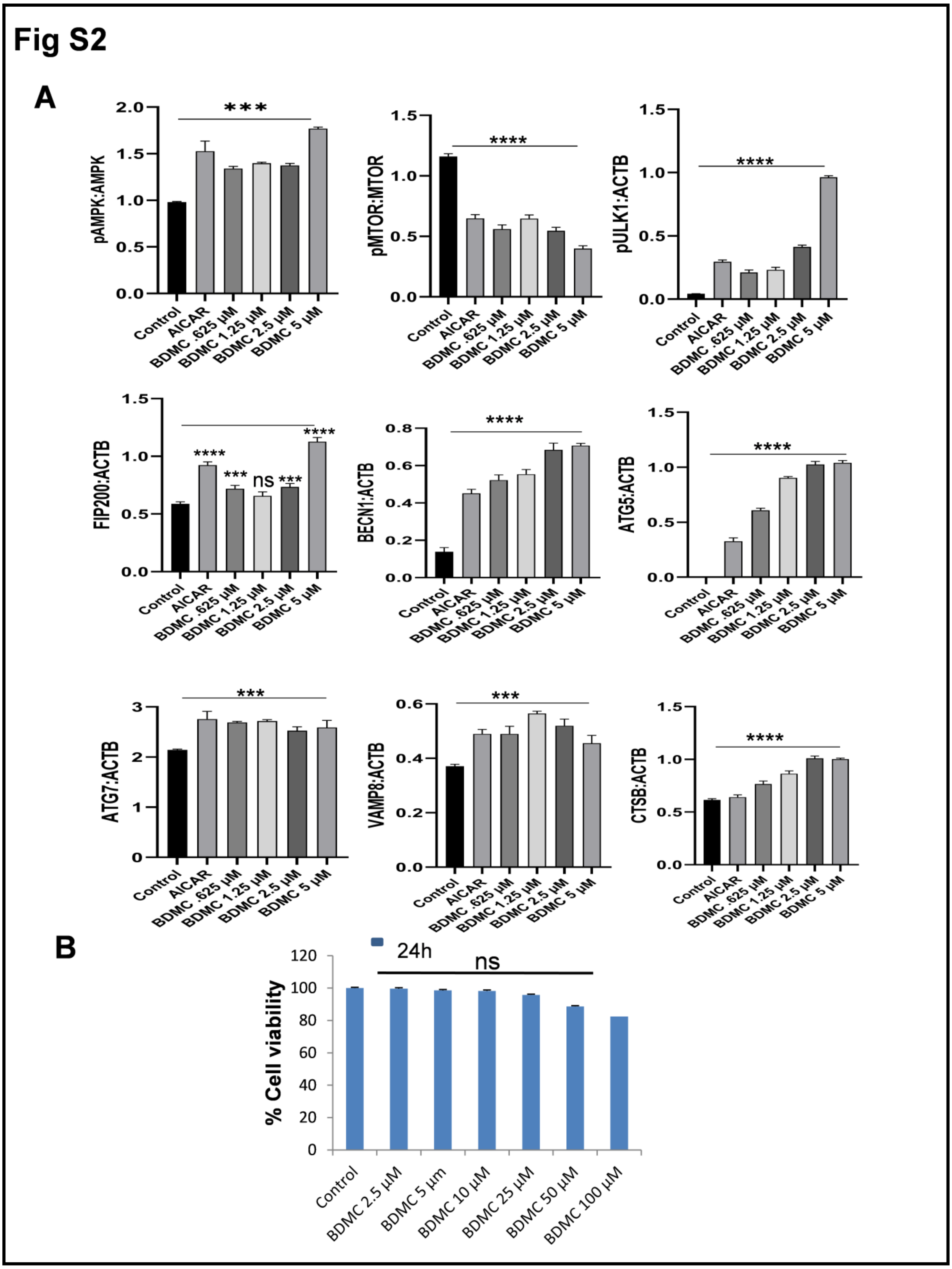


**Figure S2**: (A) Quantification of western blots shown in the Figure 2D. These data have been quantified using ImageJ software after taking densitometry values for the blots. Protein expression were normalised with ACTB and phosphorylated forms were normalised with their respective pro forms. (B) Primary astrocytes were treated with different concentrations of BDMC for 24 h to check the cell viability analysed through MTT assay. BDMC displayed no toxicity at 100 μM after treatment for 24 h in primary astrocytes. Samples were compared statistically using bonferroni test and p values $<0.05$ was considered to be significant with values ****$p < 0.0001$ ***$p < 0.001$ **$p < 0.01$

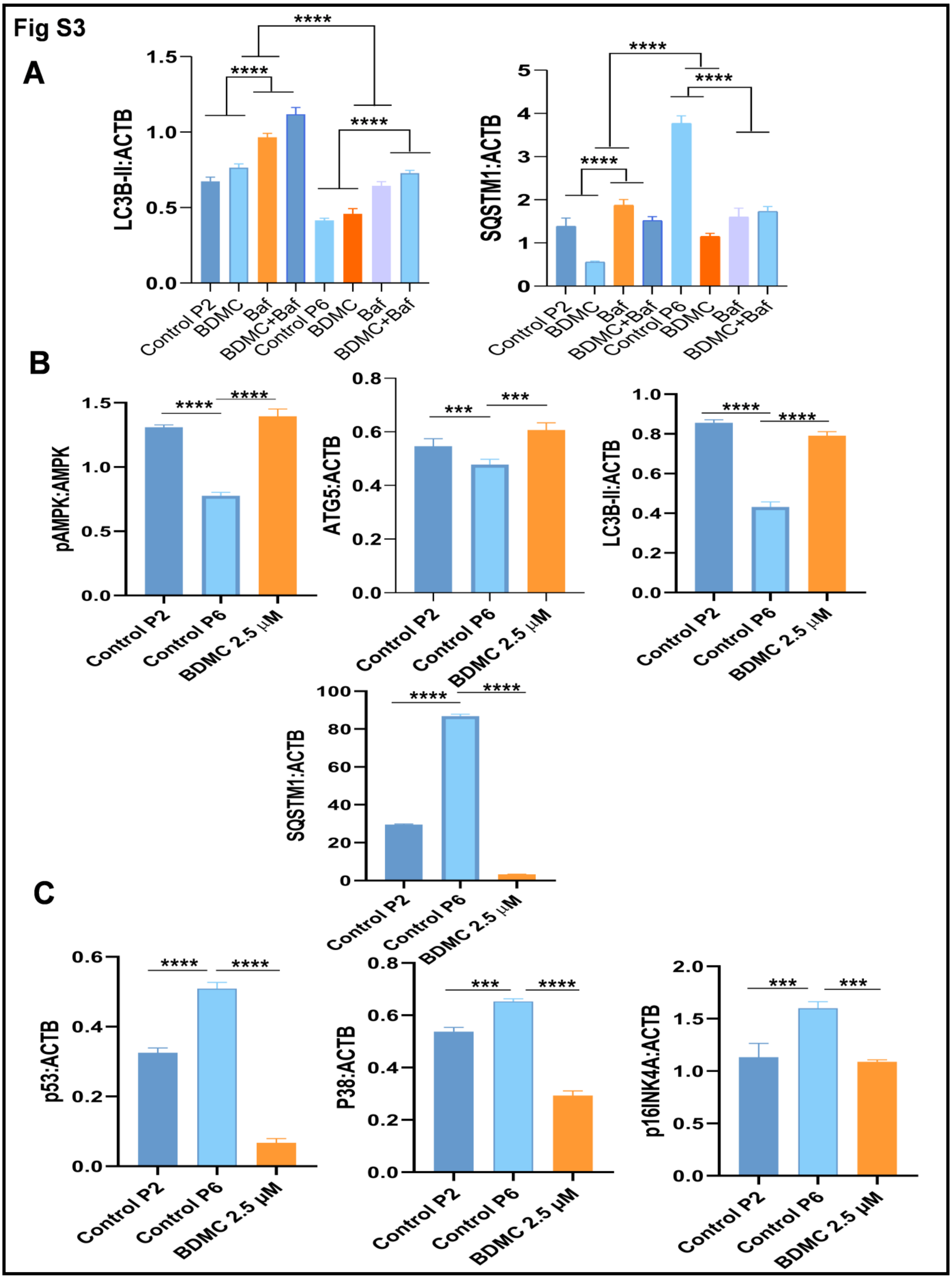


**Figure S3** (A) Quantification of western blots depicting levels of LC3B-II and SQSTM1 of Fig 3A after treatment with BDMC for 24 h in presence and absence of bafilomycin in young and aged astrocytes and normalised with ACTB. (B, C)

Quantification of expression of proteins associated with autophagy and senescence after prolonged treatment of aged astrocytes with BDMC analysed through western blotting. Samples were compared statistically using one way ANOVA and bonferroni test and p values <0.05 was considered to be significant with values ****p < 0.0001 ***p < 0.001 **p < 0.01

**Fig S4**

**A**

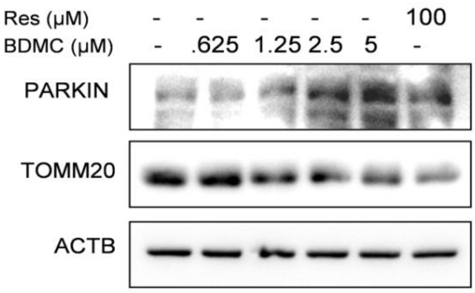


**D**

| P2 ASTROCYTES | % ROS Generation |
|---|---|
| Control | 100 |
| BDMC 2.5μM | 83.68416 |

| P6 ASTROCYTES | % ROS Generation |
|---|---|
| Control | 213.3144 |
| BDMC 2.5μM | 175.0811 |

**B**

**C**

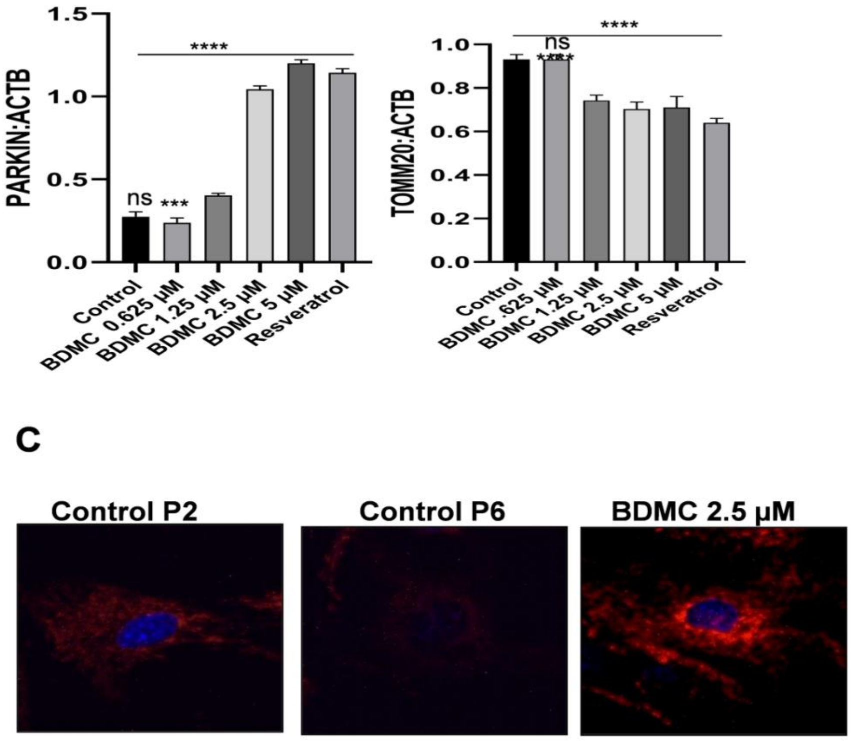

**Figure S4** (A, B) Immunoblots of mitochondria associated proteins Parkin and TOMM20 and their densitometry. Samples were compared statistically using one way ANOVA and bonferroni test and p values $<0.05$ was considered to be significant with values ****$p < 0.0001$ ***$p < 0.001$ **$p < 0.01$. (C) Representative IF images of young and aged primary astrocytes treated with BDMC and analysed for their mitochondrial activity in Fig 4A. (D) Cellular ROS generation in young and aged astrocytes. Statistically significance of the data calculated through one way ANOVA and Bonferroni analysis. p values ****$p < 0.0001$ ***$p < 0.001$ **$p < 0.01$

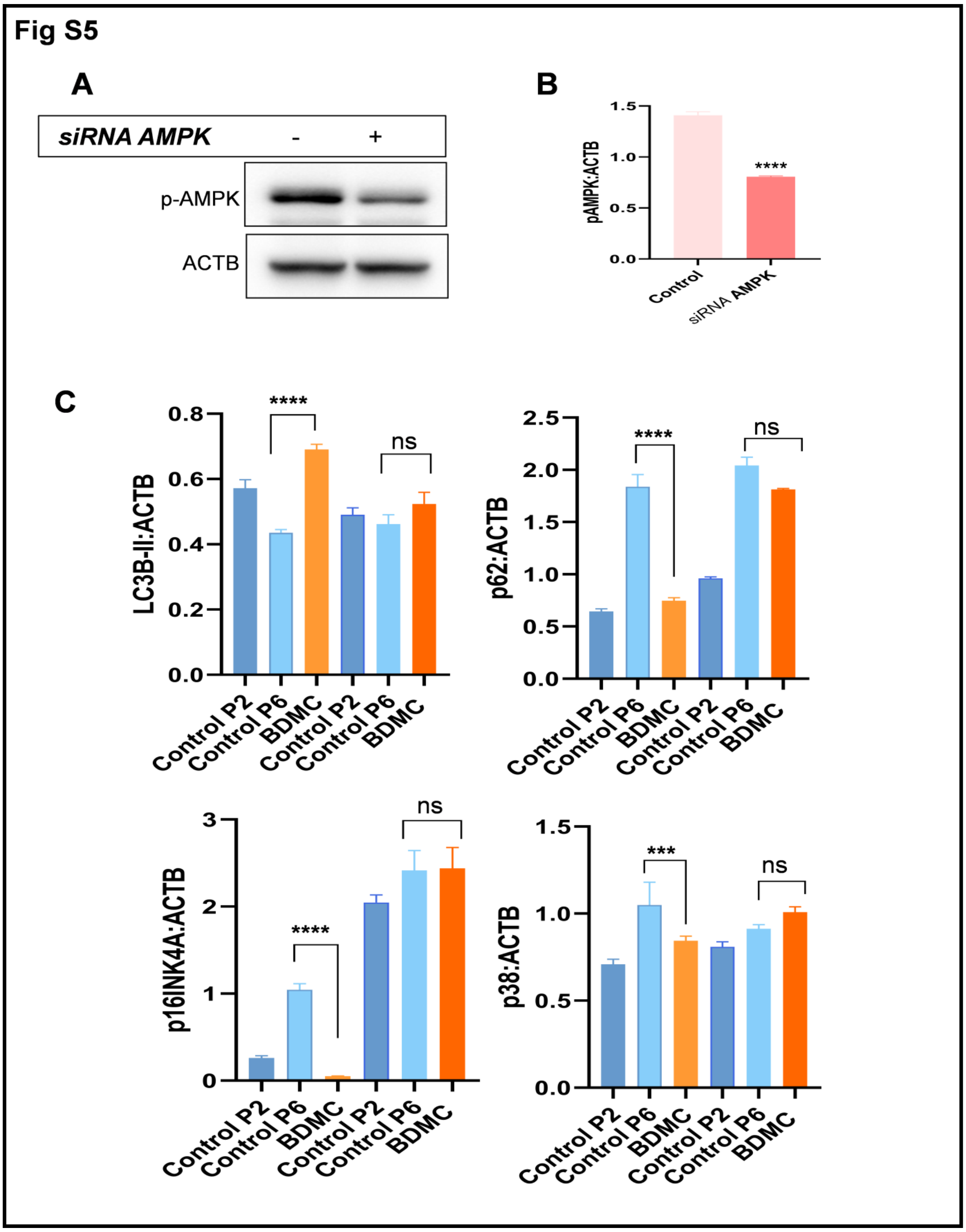


**Figure S5** (A,B) Efficacy of siAMPK was calculated by using western blot analysis and samples were normalised using value of loading control ACTB. (C) Quantification of western blots for densitometric analysis of protein levels of fig 5A. Samples were compared statistically using one way ANOVA and bonferroni test and p values <0.05 was considered to be significant with values ****p < 0.0001 ***p < 0.001 **p < 0.01

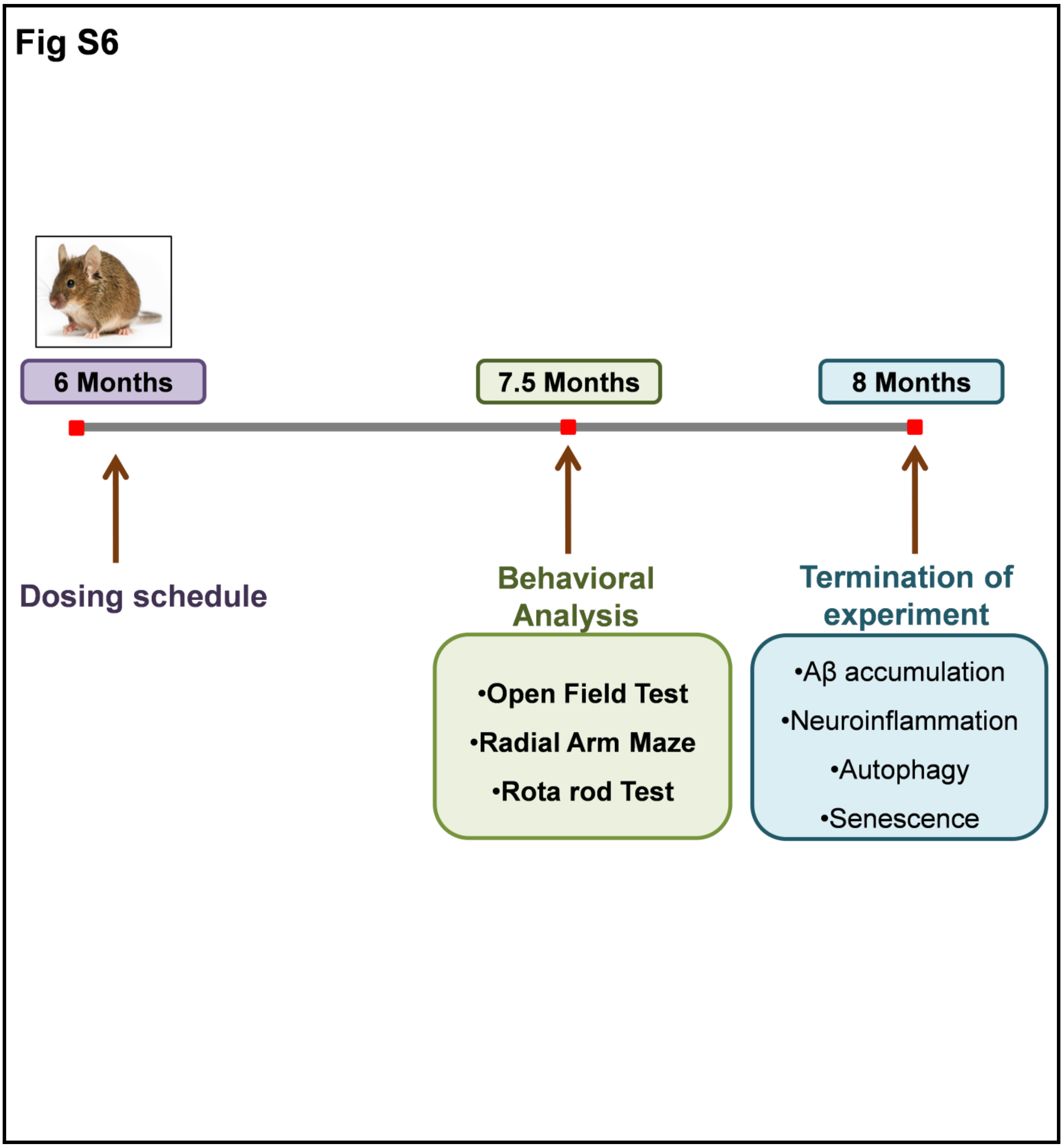


**Figure S6** Study plan for BDMC in 3xTg-AD transgenic mice. Oral dosing of BDMC was administered at 50 mg/kg and 100 mg/kg concentration to six months of 3xTg-AD mice. After 1.5 months, behavioral experiments, radial arm maze, open field test and rotarod were initiated for two weeks. The study was terminated at the end of 8$^{th}$ month by sacrificing mice and collecting brain tissue. Levels of $A\beta_{42}$ deposits were seen in the hippocampus through IHC assay. To measure the SASPs, levels of IL-1β, TNF-α and IL6 were analyzed in cortex tissue through ELISA and GFAP expression was seen in the hippocampus via IHC assay. Levels of autophagy and senescence proteins were also analyzed in the hippocampus through immunoblotting.

**Table 1**

| PK parameters | Value |
|---|---|
| $T_{1/2}$ (h) | 7.4 |
| $C_{max}$ (ng/mL) | 113 |
| $T_{max}$ (h) | 0.25 |
| $AUC_{0\text{-}t}$ (ng.h/mL) | 131 |
| $AUC_{0\text{-}\infty}$ (ng.h/mL) | 291 |
| $V_d$ (L/kg) | 1842 |
| Cl (L/h/kg) | 172 |

**Table 1** Pharmacokinetics parameters of BDMC (50 mg/kg) in C57BL/6J representing highest plasma concentration of 113 ng/ml.
